# Alpha-Gal Syndrome: Involvement of *Amblyomma americanum* α-D-galactosidase and α-1,4 Galactosyltransferase enzymes in α-gal metabolism

**DOI:** 10.1101/2021.09.13.460140

**Authors:** Surendra Raj Sharma, Gary Crispell, Ahmed Mohamed, Cameron Cox, Joshua Lange, Shailesh Choudhary, Scott P. Commins, Shahid Karim

## Abstract

Alpha-Gal Syndrome (AGS) is an IgE-mediated delayed-type hypersensitivity reaction to the oligosaccharide galactose-⍰-1,3-galactose (α-gal) injected into humans from the lone star tick (*Amblyomma americanum*) bite. This study aims at the functional characterization of two tick enzymes, α-D-galactosidase (ADGal) and α-1,4 galactosyltransferase (β-1,4GalT) in α-gal metabolism. The ADGal enzyme cleaves terminal α-galactose moieties from glycoproteins and glycolipids, whereas β-1,4GalT transfers α-galactose to a β1,4 terminal linkage acceptor sugars: GlcNAc, Glc, and Xyl in various processes of glycoconjugate synthesis. An RNA interference approach was utilized to silence ADGal and β-1,4GalT in *Am. americanum* to examine their functional role in α-gal metabolism and AGS onset. Silencing of ADGal led to the significant down regulation of genes involved in galactose metabolism and transport in *Am. americanum*. Immunoblot and N-glycan analysis of the *Am. americanum* salivary glands showed a significant reduction in ⍰-gal levels in silenced tissues. However, there was no significant difference in the level of ⍰-gal in β-1,4GalT silenced tick salivary glands. A basophil-activation test showed a decrease in the frequency of activated basophil by ADGal silenced salivary glands. These results provide an insight into the role of α-D galactosidase & β-1,4GalT in tick biology and the probable involvement in the onset of AGS.

## Introduction

Lone star ticks (*Amblyomma americanum*) transmit a variety of viral and bacterial pathogens to mammals (Childs & Paddock, 2003; Goddard & Varela-Stokes, 2009; Sayler et al., 2014). The lone star tick is also associated with southern tick-associated rash illness (STARI), and Alpha-Gal Syndrome (AGS), a newly emerged delayed allergic reaction that occurs 3-6 hours after eating beef, pork, or lamb (Commins et al., 2011; Commins & Platts-Mills, 2013). The development of specific IgE antibodies to the oligosaccharide galactose-α-1,3-galactose (α-gal) following the *Am. americanum* bites cause red-meat allergy (Commins et al., 2011; Van Nunen et al., 2009; Wuerdeman & Harrison, 2014; Crispell et al., 2019). Alpha-gal is found in the tissues of most mammals, including cattle, sheep, and swine but notably absent from humans and Great Apes. AGS is already common in several world regions; within 10 years the US alone has confirmed a spike in cases from 12 in 2009 to > 34,000 in 2019 (Binder et al., 2021) most strongly attributed to sensitization to α-gal is *Am. americanum* (Crispell et al., 2019). Several studies have also reported the presence of α-gal in the midguts of *Ixodes ricinus*, salivary glands of *Haemaphysalis longicornis, Am. sculptum,* and both the salivary glands and saliva of *Ixodes scapularis, Am. americanum* (Araujo et al., 2016; Hamesten et al., 2013a, 2013b; Chinuki et al., 2016; Crispell et al., 2019; Choudhary et al., 2021).

AGS is most common in areas where the *Am. americanum* has been historically prevalent and expanded into the regions (e.g., Long Island, NY, known for the prevalence of *Ix. Scapularis*) (Monzón et al., 2016; Sonenshine 2018). Range expansion of *Am. americanum* presents a significant public health threat in the northeastern US and beyond (Sonenshine 2018; Springer et al., 2015; Raghavan et al., 2019) due to its established role in pathogen transmission and its link to AGS (Monzón et al., 2016; Childs and Paddock, 2003; Sharma and Karim, 2021; Crispell et al., 2019, Commins, 2020). The unexpected increase in AGS is a unique health concern because strict avoidance of the food allergen is the only way to prevent a life-threatening allergic reaction.

The IgE immune response associated with food allergies is classically directed against protein antigens; however, AGS is characterized by IgE that binds to the oligosaccharide epitope galactose-α-1,3 galactose (α-gal), a cross-reactive carbohydrate determinant (CCD), specifically found in all non-primate mammals (Aalberse et al., 1981). The α-gal appears to be a common component of mammalian glycoconjugates such as glycolipids and glycoproteins. These cellular glycoconjugates are synthesized by a large family of glycosyltransferases (Berg et al., 2014; Roseman, 2001; Hennet, 2002) found throughout the cells, tissues, and fluids of all lower mammals (Apostolovic et al., 2014; Galili & Avila, 1999; Hilger et al., 2016; Iweala et al., 2020; Takahashi et al., 2014).

Our earlier combinatorial approach using N-glycome and proteome identified the α-gal antigen in salivary gland extracts and saliva of *Am. americanum* and *Ix. scapularis* (Crispell et al., 2019). When ticks feed on humans, they inject saliva, which delivers antigens containing α-gal epitopes triggering the production of anti-α-Gal antibodies (Anti-Gal) (Commins and Platts-Mills 2013). Thus, tick salivary antigens appear to be critical in the development of AGS (Crispell et al., 2019; Choudhary et al., 2021). Mysteriously, the enzyme α1,3GalT, which synthesizes α-gal, remains unidentified in tick genomes. New studies have shown indirect evidence that *Anaplasma phagocytophilum* infection in *Ixodes scapularis* cell culture induces an increase in expression of three other galactosyltransferases associated with increased levels of α- gal glycans (Cabezas-Cruz et al., 2018). However, the exact mechanism of α-gal biosynthesis in a tick is yet to be understood and, equally, the exact mechanism of how a tick bite sensitizes humans and leads to AGS development is yet to be clarified. During prolonged tick attachment on the host, the tick secretes and delivers a plethora of salivary proteins possibly containing α-gal-antigens to the host skin that might trigger an α-gal-directed IgE response (Araujo et al., 2016; Crispell et al., 2019; Choudhary et al., 2021). Surprisingly, continued exposure to ticks seems to augment the already existing IgE antibody response. However, it remains a challenge to understand why the response is so strong and directed so consistently against the α-gal carbohydrate residue. In this study, we characterized the functional role of α-D-galactosidase (ADGal) and β-1,4-galactosyltransferase (β-1,4-GalT) in galactose metabolism of *Am*.

## Materials and Methods

### Ethics statement

All animal experiments were performed in strict accordance with the recommendations in the Guide for the Care and Use of Laboratory Animals of the National Institutes of Health, USA. The protocol for tick blood-feeding on sheep was approved by the Institutional Animal Care and Use Committee (IACUC) of the University of Southern Mississippi (protocol #15101501.2). All steps were taken to alleviate animal suffering.

### Materials

All common laboratory supplies and chemicals were procured through Bio-Rad (Hercules, CA, USA), Sigma-Aldrich (St. Louis, MO, USA), and Fisher Scientific (Grand Island, NY, USA) unless specifically noted.

### Ticks and other animals

Adult unfed lone-star ticks (*Amblyomma americanum*) were purchased from Oklahoma State University’s tick rearing facility (Stillwater, OK, USA) and maintained at the University of Southern Mississippi following an established protocol (Patrick & Hair, 1975). Adult ticks were kept at room temperature at approximately 90% humidity with a photoperiod of 14 hours of light and 10 hours of darkness prior to infestation on sheep. The adult ticks were fed on sheep for time intervals between 1 and 11 days for tissue collection, depending on the experimental plan.

### DsRNA Synthesis & Tick Injections

The gene of interest was amplified using PCR with gene-specific primers and purified using the QIAquick PCR Purification Kit (QIAGEN, Germany). Gene-specific T7 promoter sequences were added to the 5’ and 3’ end of the purified product using PCR and were purified. The purified T7 PCR products were confirmed by sequencing and transcribed into dsRNA using the T7 Quick High Yield RNA Synthesis Kit (New England Biolabs, Ipswich, MA). The dsRNA produced was purified via ethanol precipitation, and the concentration was measured using a Nanodrop spectrophotometer and was analyzed on a 2% Agarose gel. Unfed females were injected with 500ng of the purified dsRNA using a 31-gauge needle and were maintained at 37°C with 90% humidity for 24 hrs. The ticks were then fed on sheep. The ticks were removed at different time points to determine the expression (Bullard et al., 2016).

### Tick tissue dissection and salivary gland extract

Partially fed female ticks removed from the sheep were dissected, and the salivary glands and midguts were removed and cleaned in ice-cold M199 buffer. Salivary glands and midguts from each time point were pooled together according to tissue type and stored in RNAlater (Life Technologies, Carlsbad NM) at -80°C until used (Bullard et al., 2016; 2019). Tick salivary protein was extracted from partially blood-fed female *Am. americanum* following the method described previously (Crispell et al., 2019). The salivary protein extracts were stored immediately at −80°C until subsequent western blot analysis.

### RNA isolation and cDNA synthesis

Frozen tick tissues were placed on ice to thaw and followed by careful removal of RNAlater. RNA was isolated from the time point pooled salivary glands using Illustra RNAspin Mini kit (GE Healthcare Lifesciences) protocols. RNA concentration was measured using a Nanodrop spectrophotometer and stored at -80°C or used immediately. Two µg of RNA was reverse transcribed using the iScript cDNA synthesis kit (Bio-Rad) to synthesize cDNA. The reverse transcription reaction is then heated in a Bio-Rad thermocycler under the following conditions: 5 minutes at 25°C, 30 minutes at 42°C, 5 minutes at 85°C, and hold at 10°C. The resultant cDNA was diluted to a working concentration of 25 ng/µl with nuclease-free water and stored at -20°C until used (Bullard et al., 2016).

### Quantitative Real-Time PCR

QRT-PCR was performed within the guidelines of Bio-Rad protocols provided with iTaq Universal SYBR Green Supermix. Briefly, 50 ng of cDNA was added to a 20 µl qRT-PCR reaction using SYBR Green supermix with 300 nM of each gene-specific primer. The samples were subjected to the following thermocycling conditions: 95°C for 30 sec; 35 cycles of 95°C for 5 sec and 60°C for 30 sec with a fluorescence reading after each cycle; followed by a melt curve from 65°C to 95°C in 0.5°C increments. Each reaction was performed in triplicate along with no template controls (Bullard et al., 2016). Primers used for gene expression validation can be found in Supplementary Table 1. Gene expression validation was performed using β-actin and histone as the reference gene.

### Quantification of total bacterial load

The total bacterial load in tick tissues was determined using the method described elsewhere (Budachetri and Karim 2015; Narasimhan et al., 2014). Briefly, 25 µl volume reaction mixture contained 25 ng of tissue cDNA, 200 µM 16S RNA gene primer, and iTaq Universal SYBR Green Supermix (Bio-Rad) followed by a qPCR assay using following conditions: 94 °C for 5 min followed by 35 cycles at 94°C for the 30s, 60°C for 30s and 72°C for 30s. A standard curve was used to determine the copy number of each gene. The bacterial copy number was normalized against *Am. americanum* actin copy number in control tissues and gene silenced tick, and each sample was run in triplicate.

### SDS-PAGE and Immunoblotting

SDS-Polyacrylamide Gel Electrophoresis and Immunoblotting were carried out using the methods described elsewhere (Crispell et al., 2019). Proteins extracted from the salivary glands (15 µg) were fractionated on a Mini-PROTEAN TGX Any kD, 4–20% gel (Bio-Rad) using SDS-PAGE. They were then transferred onto a nitrocellulose membrane in a Transblot cell (Bio-Rad). The transfer buffer consisted of 25 mM Tris-HCl and 192 mM glycine in 20% methanol. Blocking of nonspecific protein binding sites was executed with 5% BSA in a TBS and Tween-20 solution. The membranes were incubated with α-galactose (M86) monoclonal IgM antibodies (Enzo Life Sciences, Farmingdale, NY, USA) at a dilution of 1:10 using an iBind western device (Life Technologies, Camarillo, CA, USA). The antigen-antibody complexes were visualized using a secondary horseradish peroxidase-conjugated goat anti-mouse IgM antibody (Sigma-Aldrich) at a dilution of 1:10,000. They were detected with SuperSignal chemiluminescent substrate (Pierce Biotechnology, Rockford, IL, USA) using a Bio-Rad ChemiDox XRS.

### Basophil Activation Assay with Tick Salivary glands

Peripheral blood mononuclear cells (PBMCs) taken from a healthy, non-α-gal allergic donor (α-gal sIgE <0.10) were isolated using a Ficoll–Paque gradient (GE Healthcare, Chicago, IL, USA). Endogenous IgE was stripped from basophils within the PBMC fraction by incubating the cells with cold lactic acid buffer (13.4 mM lactic acid, 140 mM NaCl, 5 mM KCl) for 15 min. Basophils were sensitized with plasma from α-gal allergic and non-allergic subjects overnight in RPMI 1,640 cell culture media (Corning CellGro, Manassas, VA, USA) in the presence of IL-3 (1 ng/mL, R&D Systems, Minneapolis, MN, USA) at 37°C and 5% CO2. PBMCs were subsequently stimulated for 30 min with RPMI media, cetuximab (10 μg), rabbit anti-human IgE (1 μg; Bethyl Laboratories Inc., Montgomery, TX, USA), partially fed salivary gland extracts from *Am. americanum* (50 μg). Stimulation reactions were stopped with 20 mM EDTA, and PBMCs stained with fluorescently labeled antibodies against CD123 (BioLegend, San Diego, CA, USA), human lineage 1 (CD3, CD14, CD16, CD19, CD20, CD56, BD Biosciences, San Jose, CA, USA), HLA-DR, CD63 (eBiosciences, ThermoFisher, Waltham, MA, USA), and CD203c (IOTest Beckman Coulter, Marseille, France) in flow cytometry staining buffer with 0.02% NaN3. Samples were acquired on a CyAN ADP flow cytometer (Beckman Coulter, Brea, CA, USA) and analyzed using FlowJo v10 software (FlowJo LLC, Ashland, OR, USA). Data analysis was performed using Prism version 7.03 (GraphPad Software, La Jolla, CA, USA). Mann–Whitney U-tests were used to compare the frequency of CD63+ basophils detected following stimulation with various compounds. A p-value < 0.05 was considered significant.

### N-Glycome analysis of tick salivary glands

N-linked glycans were released from 30 μL of *Am. americanum* salivary glands with an estimated protein concentration of 200 μg, after being reduced, alkylated, and then digested with trypsin in Tris-HCl buffer overnight. After protease digestion, the sample was passed through a C18 seppak cartridge, washed with 5% v/v acetic acid, and the glycopeptides were eluted with a blend of isopropanol in 5% v/v acetic acid before being dried by SpeedVac. The dried glycopeptide eluate was treated with a combination of PNGase A (Sigma) and PNGase F (New England Biolabs, Ipswitch, MA, USA) to release the N-linked glycans. The digest was then passed through a C18 sep pak cartridge to recover the N-glycans. The N-linked glycans were then permethylated for structural characterization by mass spectrometry. Briefly, the dried eluate was dissolved with dimethyl sulfoxide and methylated with NaOH and methyl iodide. The reaction was quenched with water, and per-O-methylated carbohydrates were extracted with methylene chloride and dried under N2. The permethylated glycans were reconstituted in 1:1 MeOH: H_2_O containing one mM NaOH, then introduced to the mass spectrometer (Thermo Fusion Tribrid Orbitrap) with direct infusion at a flow rate of 0.5 μL/min. Full MS spectra and an automated “TopN" MS/MS program of the top 300 peaks were collected and fragmented with collision-induced fragmentation. These fragmentation data were used to confirm a Hex-Hex-HexNAc signature, both with a diagnostic fragment, as well as expected neutral losses.

## Results

### Temporal expression of α-D-galactosidase and β-1,4-galactosyltransferase in tick salivary glands

Temporal expression in *Am. americanum* salivary glands revealed the α-D-galactosidase expression level increases ∼2-fold after tick attachment to the host during the slow feeding phase up to five days post-infestation (dpi), and it decreased ∼2-fold at seven dpi and ten dpi, during the rapid feeding phase (Figure 1A). Interestingly, β-1,4- galactosyltransferase transcript level increases ∼8-fold after attachment at three dpi and remains upregulated (∼2 fold at 5dpi; ∼4.5 fold at seven dpi and ∼2 fold at ten dpi) compared to the unfed salivary glands. Transcriptional expression was normalized against the unfed tick salivary gland. β-actin and histone, two housekeeping genes, were used to normalize the gene expression.

**Figure 1:**
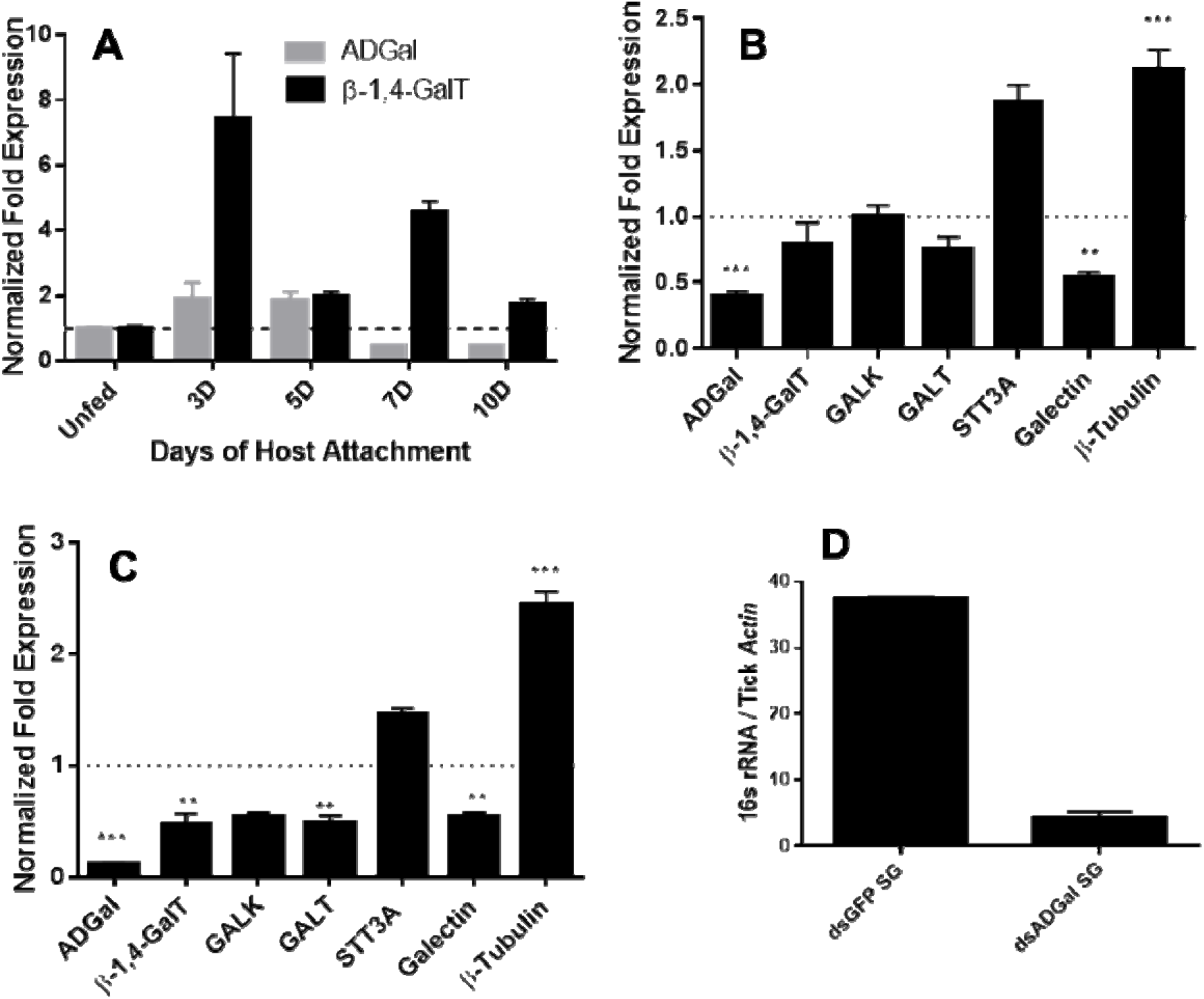
Transcriptional expression in tick tixssues. **A)** time-dependent transcriptional gene expression of α-D-galactosidase (ADGal) and β- 1,4 galactosyltransferase(β- 1,4-GalT) in *Amblyomma americanum* salivary glands. Fold changes were normalized with the unfed tissue expression. Β-actin and histone were used to normalize the gene expression. **B)** Transcriptional expression of carbohydrate metabolism and transport-related genes in α-D-galactosidase silenced partially blood752 fed midguts and, **C)** salivary glands. Β-actin and histone, house-keeping genes were used to normalize the expression against dsGFP treated tissues. Abbreviations: β- 1,4- GalT ( β-1,4 galactosyltransferase); GALK (Galactokinase); GALT (galactosyltransferase); STT3A (Dolichyl-diphosphooligosaccharide-protein glycosyltransferase)(*P<0.005; **P<0.01,***P<0.001, student t-test), **D)** Total bacterial load partially blood-fed salivary glands injected with dsGFP and dsADGal. Total bacterial load was quantified by qPCR using β-actin as a reference gene.

### Gene silencing and transcription expression of genes related to galactose metabolism

α-D-galactosidase dsRNA injections led to ∼65% and ∼85 % downregulation of ADGal gene expression in both midguts and salivary glands (Figure 1B-1C), respectively. Transcriptional expression analysis of α actosidase silenced tick tissues shows the significant downregulation of galectin (∼50%), while a significant upregulation of β tubulin in the midgut (∼2-fold increase) and salivary gland tissues (∼2.5-fold increase). In the salivary glands, a significant downregulation of β 4-GalT of approximately 2-fold and GALT by approximately 2-fold, and a non-significant decrease in galactokinase (GALK) were noted. In addition to that, there was upregulation of STT3A gene in ADGal silenced midgut (∼1.6-fold increase) and salivary gland (∼0.5-fold increase); however, this change was not statistically significant. Furthermore, there was non-significant downregulation of β 4-GT, GALT in ADGal silenced midgut tissues.

### α-D-galactosidase silencing on Bacterial load

Total bacterial load quantification assay showed that ∼7-fold reduced 16S bacterial load in the salivary gland tissues of *Am. americanum* ticks that received dsADGal injections, compared to dsGFP irrelevant control injected ticks (Figure 1D).

### Impact of gene silencing on feeding success and tick engorgement

To investigate the impact of silencing of α-galactosidase and β-1,4- galactosyltransferase on the tick phenotype, we measured and compared engorged tick weight. Ticks treated with dsADGal engorged faster and weighed more than ticks injected with dsGFP irrelevant control (Figure 2). The dsAGS tick weights were significantly (P<0.05) increased at ten dpi compared with dsGFP control ticks. However, there was no significant difference in tick engorgement in β-1,4-galactosyltransferase gene silenced ticks.

**Figure 2.**
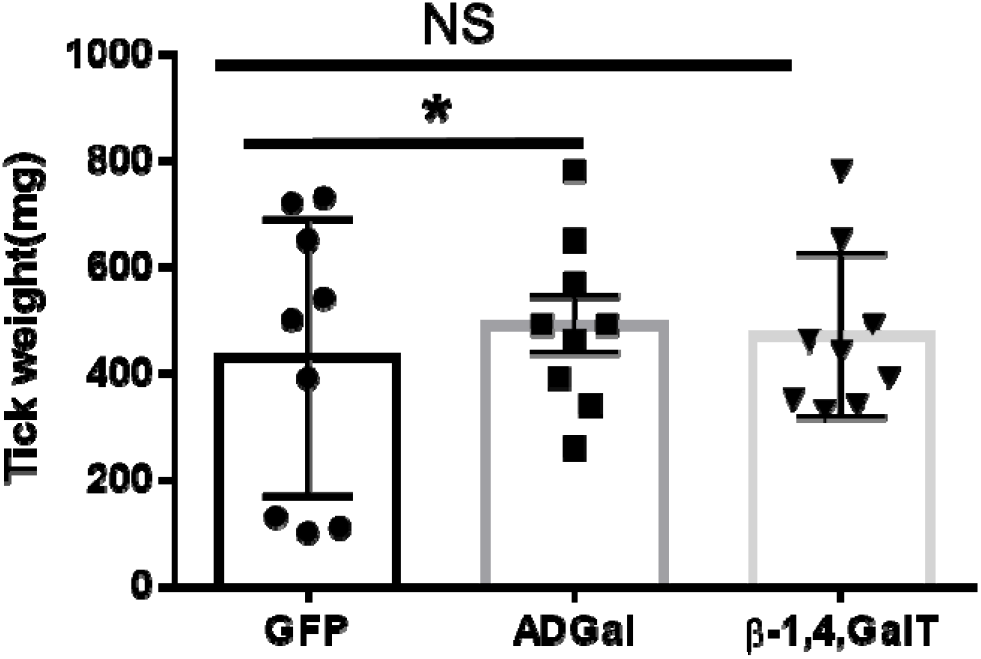
Engorgement weights of ticks treated with dsRNA, tick weights were taken from mass replete or forcibly removed ticks at four time-points during the bloodmeal after treatment with dsADGal (α-galactosidase), dsβ- 1,4-GT(β-1,4 galactosyltransferase) or dsGFP (irrelevant control) double-stranded RNA (* P<0.05, students t-test )

### Alpha-gal expression in gene silenced tissues

Salivary glands from five dpi, seven dpi, and nine dpi *Am. americanum* ticks injected with dsADGAL and dsGFP irrelevant control RNA were assessed using immunoblotting with an anti-alpha-gal IgM antibody (Figure 3). Densitometry analysis was conducted to determine the relative abundance of α in dsADGal injected tick protein against dsGFP control protein (Supplementary figure 3). Results indicate an ∼80% reduction in α-gal in dsADGal salivary gland proteins compared to the dsGFP control. A decrease in α-gal of more than 30% in the seven dpi dsADGal injected tick salivary glands, but the nine dpi salivary glands contained ∼10% more α than the dsGFP irrelevant control. While there was no significant difference in α ds β 4-GT injected *Am. americanum* ticks salivary gland in comparison to control (Supplementary figure 2B).

**Figure 3.**
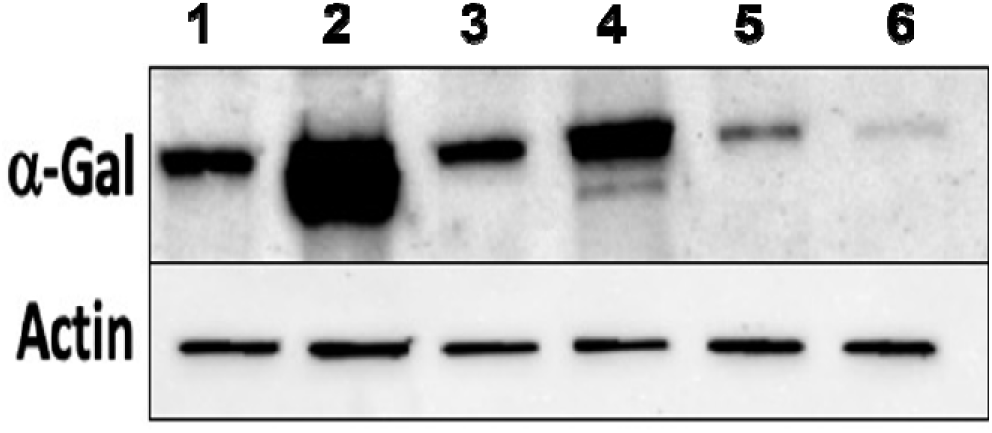
Western blot analysis of α-gal using anti-gal IgM in dsADGal (α-galactosidase) and dsGFP(irrelevant control) injected partially-fed *Am. americanum* salivary glands. Lane 1, 5days post infestation (dpi.) dsADGal salivary glands; Lane 2, 5dpi dsGFP salivary glands; Lane 3. 7dpi dsADGal salivary glands; Lane 4. 7dpi dsGFP salivary glands; Lane 5. 9dpi dsADGal salivary glands; Lane 6, 9dpi dsGFP salivary glands

### Silencing of α-D-actosidase reduces α-D-containing cross-reactive carbohydrate determinants (CCDs) in tick salivary glands

We performed N-glycan analysis to check the impact of α-D-actosidase silencing on the abundance of α containing glycoforms in tick salivary glands. Profile analysis of N-glycan from control samples and dsADGal showed a variety of high mannose, complex type, fucosylated, and alpha-gal containing glycoforms (Table 1, Supplementary figure 3-4; Supplementary tables1-3) which were similar in overall trends to the N-glycan profile published previously (Crispell et al., 2019). In addition, the overall α glycoforms abundance profile showed that, in both samples, the abundance of fucosylated glycoforms was higher than non-fucosylated glycoforms (Supplementary tables: 1-3; Supplementary figures: 3-4). More specifically, overall N-glycans abundance containing α gal glycoforms or moieties in dsGFP injected control tick salivary gland was 24.02%; however, in the dsADGal treated salivary glands, overall N-glycans containing α moieties were significantly reduced to 2.81%. Among these data, the α having glycoforms at m/z of 2478 and 2723 were absent in the dsADGal treated salivary glands while other glycoforms m/z 2652 and 2897 compared to control (Table 1, Supplementary tables: 1-3). These results strengthen the hypothesis that tick α-D-galactosidase is vital in synthesizing or transfer of α-gal to tick salivary glycoproteins.

**Table 1:** N-Glycan analysis on 5dii *Am. americanum* salivary gland protein extracts. The mass to charge ratio (m/z) signifies different α-gal glycoforms. Total percentage α- gal indicates the percentage sum of different α-gal glycoforms detected in SG-Control dsGFP or irrelevant control) treated and ds α-galactosidase-treated (SG-ADGal KD).Greyed out Boxes indicate that this mass was not detected in that sample. Key: NO: MS/MS fragmentation resulted in 486.23 ion

|  |  |  |  |  |  |  |  |  | TOTAL<br>%<br>Alpha-Gal |
| --- | --- | --- | --- | --- | --- | --- | --- | --- | --- |
| m/z | 1825 | 1999 | 2070 | 2244 | 2478 | 2652 | 2723 | 2897 |  |
| SG-Control |  | 0.25% | NO | 0.00% | 2.10% | 8.94% | 4.43% | 8.30% | 24.02% |
| SG-KD | 0.11% | 0.51% | NO | 0.41% |  | 1.23% |  | 0.46% | 2.81% |

Galactose N-acetylglucosamine Mannose Hexose

**Table 1:**
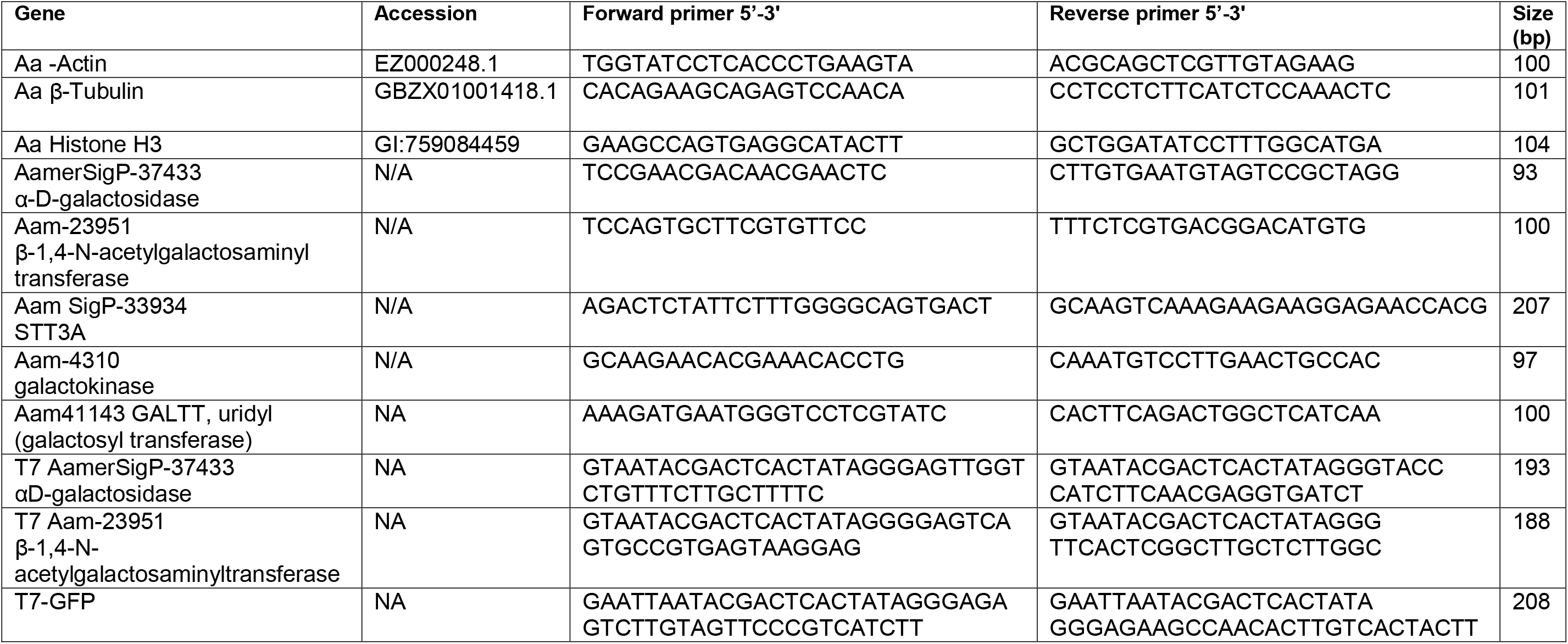
List of genes, accession numbers, primers, and base sizes used in this study for transcriptional and gene silencing experiments.

### Basophil Activation Test with tick salivary gland samples

Since the profile of N-glycan demonstrates the presence of α-gal antigen in salivary samples, we sought to analyze the impact of ADGal silencing in basophil activation. In this basophil activation test, the frequency of CD63+-activated donor basophils is lower when PBMCs are stimulated with ADGal silenced five dpi *Am. americanum* salivary gland protein extract in comparison to control five dpi *Am. americanum* salivary glands, cetuximab, and Anti-IgE positive control (Figure 4). More specifically, we found that the frequency of CD63+ basophils was significantly increased following sensitization with α gal allergic plasma and stimulation with α-D-containing tick salivary extract samples from *Am. americanum* (PF SG extract) (p < 0.05 vs. media, 4).

**Figure 4:**
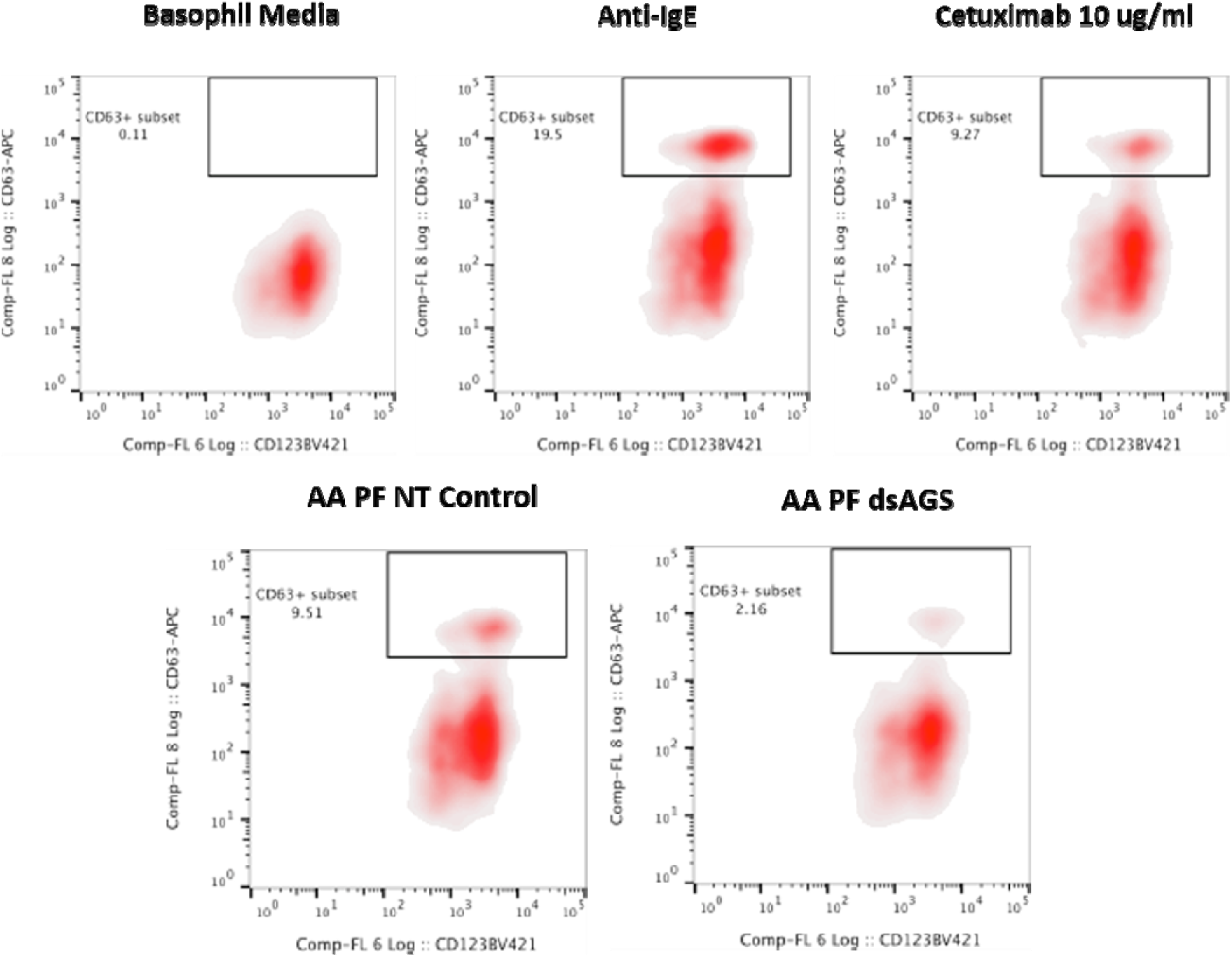
Flow cytometric analysis of human basophil activation by *Amblyomma americanum* salivary proteins. Donor basophils from a healthy, non-allergic control subject were stripped of IgE and primed overnight with plasma from a subject with α-gal syndrome (α-gal sIgE = 31.3 IU/mL, total IgE = 233 IU/mL). Sensitized cells were exposed to one of the following stimuli for 30 min: RPMI media, crosslinking anti-IgE antibody (Positive control), α-gal containing glycoprotein, cetuximab (α-gal positive control), partially blood-fed *Am. americanum* control and α-D-galactosidase gene silenced salivary gland extracts treated, CD63 expression on lineage-HLA-DR-CD123+CD203c+basophils were assessed by flow cytometry.

Furthermore, ADGal silenced salivary extract five dpi samples from *Am. americanum* showed a significantly lower frequency of CD63+ basophil (2.16%) activation. In contrast, the frequency of CD63+ activated basophils following stimulation with partially fed five dpi salivary extract was 9.51% when the results of all experiments (n=3) were combined. Overall, these results suggest a correlation between tick ADGal depletion and potential reduction in the host allergic immune response.

## Discussion

The discovery of α-gal immunoglobulin E (IgE), a central player involved in allergic responses against food containing α-gal antigen such as red meat, and in people with a history of tick bite has caught enormous attention among immunologists and vector biologists (Commins, 2020; Crispell et al., 2019; van Nunen 2009; Sharma and Karim, 2021). Several research reports have established that antigen-containing α-gal is a key trigger for AGS development (Commins, 2020). Current research focuses on identifying and profiling α-gal antigens in tick saliva and tissues to decipher the interplay between a tick bite and AGS (Sharma and Karim, 2021; Crispell et al., 2019; Cabezas-Cruz et al., 2019). It is still puzzling how the tick acquires and decorates its saliva antigens with α- gal and primes the host immune response. The expression of α-gal antigen in tick saliva is established in different ticks, including *Am. americanum*. However, answer to the questions relating to origin or source of α–gal in a tick and the key mechanistic details of how bite from this tick leads to sensitization of humans against red meat allergy either by triggering the development of memory cell capable of producing α-gal IgE or class switching of IgE because of salivary factor is yet to be clarified. Several pieces of evidence support that the presentation of cross-reactive carbohydrates determinants or the α antigen by ticks during tick feeding is possibly the prime factor for sensitization of humans against α (Choudhary et al., 2021; Crispell et al., 2019). However, how ticks acquire or produce α moiety during feeding remains a complete mystery (Hamsten et al., 2013; Arujo et al., 2016). There are few working hypotheses regarding the source of α in a tick; (a) α present in a tick is residual or enzymatically modified/ cleaved mammalian α-gal containing glycoprotein or glycolipids derived from the mammalian host, (b) α–gal is originating from tick hosted microbes (tick microbiome) capable of expressing α–gal in or possessing the capability to capture cleaved galactose oligosaccharide and glycosylate their own or tick salivary proteins (Sharma and Karim 2021).

Recently, we have demonstrated (Crispell et al., 2019; Choudhary et al., 2021; Sharma and Karim 2021): 1) tick bites, specifically by the lone star tick, might be solely responsible for stimulating an IgE response to α-gal within the southern and eastern United States. 2) N-linked glycan analysis confirmed the presence of α-gal in the saliva and salivary glands of *Amblyomma americanum* and *Ixodes scapularis*, but *Amblyomma maculatum* contained no detectable quantity. 3) An immuno-proteome approach confirmed the cross-reactivity between tick saliva proteins (allergens) to α-gal antibodies. 4) The presence of antigenic galactose-α-1, 3-galactose (α-gal) epitope activated human basophils as measured by increased expression of CD63. 5) The lone star tick salivary gland extracts induced AGS in α-Gal^-KO^ mice. An immunoproteome and sialotranscriptome analysis identified several expressed molecules during feeding progression linked with galactose metabolism, N-glycan synthesis, galactoside transport (Crispell et al., 2019; Karim and Ribeiro 2015). Surprisingly, the key enzyme, α-1,3- galactosyltransferase (α1,3GalT), synthesized α-gal, remains unidentified in tick genomes and transcriptomes. However, this enzyme is reported to be the key enzyme involved in the synthesis of α1,3-galactose (α-1,3Gal or α-gal) epitopes in several organisms (Galili, 1999; 2005; 2015).

We investigated the role of differentially expressed tick molecules during prolonged blood-feeding on the host, i.e., α-D-galactosidase (glycoside hydrolase/ADGal); an enzyme that catalyzes the breakdown of galactose from glycoproteins or glycolipids, and β-1,4-galactosyltransferase (β-1,4-GalT); an enzyme that is involved in the synthesis of Galβ1-4-GlcNac-disaccharide unit of glycoconjugates (Hennet, 2002; Calhoun et al., 1986). We hypothesized that the expression of these genes during prolonged feeding and can impact on overall α–gal expression, expression of key genes of galactose catabolism (Leloir pathway), N-glycan synthesis, galectin (a molecule involved in transport of galactose containing oligosaccharides) (Diaz and Ortega, 2017; Huang et al., 2007), and β-tubulin (a glycosylated marker molecule). Leloir pathway (Supplementary figure 1) is the predominant route of galactose metabolism and products of this pathway generate key energy molecules as substrate, i.e., UDP-galactose vital for N-glycan synthesis (Freeze et al., 2017; Karim and Ribeiro, 2015). More specifically, selected key molecules of the Leloir pathway included in this study were galactokinase (GALK), galactose-1-phosphate uridyltransferase (GALT). GALK modifies galactose to create a molecule called galactose-1- phosphate, which can be further added to build galactose-containing proteins or fats (McAuley et al., 2016). Whereas GALT catalyzes the second step of the Leloir pathway (Supplementary figure 1), converting galactose into glucose (Wong & Frey, 1974). In addition to that, another molecule, AamSigP-24522 putative dolichyl-diphosphooligosaccharide protein glycosyltransferase (STT3A), which is reported to be involved in early stage of cotranslational N-glycosylation of target protein (Cherepanova & Gilmore, 2016), was also included in the study to check the impact of silencing of ADGal and β-1,4-GalT in the initial step of N-glycosylation.

Furthermore, in this study, the impact of ADGal and β-1, 4-GalT silencing on glycosylated molecule and molecule involved in transport, the expression of β-tubulin and galectin was tested. In addition to that, in this study, we analyzed the expression of α-gal antigen in salivary glands of *Am. americanum* across the different feeding stages, we observed that α-gal antigens in partially blood-fed are significantly up-regulated (3-5 days) and gradually down-regulated as feeding transitions towards the fast feeding stage and repletion. Such concurring trend of differential expression of ADGal and α-gal antigen expression in a tick salivary gland across feeding stages points towards the role of sialome-switch in tick’s α-gal signature (Karim and Ribeiro 2015). However, the central question regarding tick’s inherent ability to express α-gal remains enigmatic because of the lack of α-1,3 galactosyltransferase sequence in tick databases. To date, there are 1,776 genomes from vertebrates available at the NCBI genome site (https://www.ncbi.nlm.nih.gov/genome/?term=vertebrata) and 2,513 arthropod genomes, 11 of which are from ticks. Likewise, the silencing of the ADGal gene in both midgut and salivary gland tissues showed an interesting compensatory expression pattern of genes involved in galactose metabolism (Supplementary fig. 1). The transcript levels of β-1, 4-galactosyltransferase, galactose-1-phosphate uridyltransferase (GALT), and galactose binding transport protein galectin were significantly down-regulated; while, the transcript level of β-tubulin was significantly upregulated. GALK, GALT, and galectin downregulation may be negative feedback responses due to the reduction of galactose-containing oligosaccharides caused by silencing of ADGal. While putative dolichyl-diphosphooligosaccharide protein glycosyltransferase subunit STT3A gene was up-regulated, upregulation was not statistically significant, probably due to compensatory effect from the redundant molecule. Since the temporal expression pattern of β-1, 4-GalT coincided with α-gal antigen expression, we also carried out a functional study using RNAi.

Regardless of significant silencing of β-1,4-GT gene in salivary gland tissues, the transcript level of ADGal, GALK, GALT, STT3A, Galectin, β-tubulin as well as α-gal- expression in *Am. americanum* showed no significant change. These results suggest β- 1, 4-GalT does not contribute to the α-gal signature in *Am. americanum*. Intriguingly, three variants of galactosyltransferase genes, i.e., β4galt-7; α4galt-1, and α4galt-2 from β-1, 4-GalT, and α-1, 4-GalT family are shown to be involved in the α-gal synthesis pathway (Cabezas-Cruz et al., 2018). However, search for *Ix. scapularis* homologs in *Am. americanum* failed to yield any CDS in the existing NCBI transcriptomic database. Since the genome sequence of *Am., americanum* is not available so far. Hence, it will be challenging to conclude the absence of such important genes.

Since ADGal is an enzyme responsible for releasing α-gal from substrates, it is differentially expressed across the feeding stages, indicating tick may be utilizing it for removing α-gal from host-derived glycosylated lipid or proteins. To answer the question that α-D-galactosidase plays a critical role in galactose metabolism and α-gal signature in the tick tissues. We silenced the ADGal gene and performed N-Glycan analysis. The results indicated a significant reduction in the overall abundance of N-glycans containing α-gal glycoforms in tick samples. Basophil activation test using control and ADGal Salivary Gland Extract (SGE) showed a significant reduction in the frequency of activated basophil compared to control. These results further confirmed the N-Glycan analysis in the gene-silenced salivary glands. The engorgement weights of dsADGal ticks compared to control ticks showed a significant difference, and gene-silenced ticks gained weight faster than control ticks. This phenotype could result from the tick’s compensatory mechanisms to losing a key galactose metabolizing molecule. The transcriptional expression data suggest the silencing of ADGal inhibits the tick’s Leloir galactose metabolism pathway by downregulating the expression of intermediate enzymes, GALK and GALT, and correlates with differential expression of other galactose-modifying genes including, β1-4, galactosyltransferase, galectin. The Leloir pathway ultimately channels into glycolysis and produces ATP. The ATP was not quantified in these experiments. Still, the downregulation of Leloir pathway genes and decrease in α-gal may have led to the tick producing less energy and compensating it by imbibing high-quantity of blood meal (Raven & Johnson, 1995). Our findings support the functional role of ADGal in the tick’s energy utilization.

A 6-fold decreased total bacterial loading in ADGal silenced partially-blood fed tick tissues supports the hypothesis that free galactose or glucose reduces the total bacterial load within the ticks. These results also support the functional role of ADGal in maintaining microbial homeostasis within the tick salivary glands. These findings further warrant investigations to examine the role of bacterial communities in AGS because of their possible role in manipulating ticks’ metabolic activity and glycosylation machinery. Microbiome homeostasis within the tick is critical in the context of the α-gal syndrome (AGS). Galactose is vital for microbes not only as an energy molecule but also as a key molecule required to produce glycosylated exopolysaccharides or lipopolysaccharides (LPS), a potential α-gal antigen (Chai et al., 2012). In addition, the presence of specific microbes in the tick vector can also affect metabolome, especially galactose. One recent study of tick- *Borrelia* interaction found that relative abundance of galactose was significantly reduced in *Borrelia burgdorferi* and *Borrelia mayonni* infected ticks (Hoxmeier et al., 2017). Likewise, an earlier report on the role of galactose and Leioler pathway genes, especially galactosyltransferase, established that galactose and bacterial galactosyl transferases are vital in biofilm formation for colonization of bacteria (Chai et al., 2012). Hamadeh et al. (1996) demonstrated glycosylation of human erythrocytes (RBCs) *in vivo* by a bacterial α1,3 galactosyl transferase enzyme. Another recent study demonstrated that the tick-borne pathogen *Anaplasma phagocytophilum* increases α-gal antigen in IRE *Ix. ricinus* tick cells (Cabezas-Cruz et al., 2016). Montassier et al. (2019) reported the presence of α1,3-galactosyltransferase bacterial sequences in the human gut microbiome shotgun sequencing project (Montassier et al., 2019). The microbes from Rizobiaceae and Caulobacteriaceae families were found to possess a novel lipid A a- (1,1)-GalA transferase gene(rgtF) (Brown et al., 2013). These findings provided a supporting basis for the hypothesis that a glycosylated lipid could be one augmenting factor for sensitization. Bacteria utilize this machinery to synthesize exogenous lipopolysaccharides (LPS), hence lipid A a-(1,1)-GalA transferase, which could be necessary for an α-gal antigen development (Brown et al., 2013; Del Moral & Martínez- Naves, 2017). Considering all these facts, it is inferred that the microbiome could also be one factor involved in the sensitization against α-gal while the tick is feeding on the host.

## Conclusion

The results from multiple research papers have led to the conclusion that the (a) tick α- D-galactosidase is an important enzyme that is uniquely expressed in salivary glands, and tick utilizes this enzyme to cleave α-gal from host proteins or lipids and recycle or add in its proteins during hematophagy (b) α-D-galactosidase silencing reduces N glycan signature (α-gal moieties) in the tick salivary glands, (c) β-1,4- galactosyltransferase downregulates galactose catabolism however silencing in a tick does not affect overall α-gal expression, (d) α-D-galactosidase silencing also significantly reduces the microbial load in tick salivary glands. Overall, α-D-galactosidase is an essential enzyme involved in the development of α-gal antigen in ticks during hematophagy. The results presented here add new insights into understanding the role of vital tick intrinsic factors involved in the synthesis or recycling of α-gal and sensitize host against α-gal during hematophagy.

## Ethics Statement

All animal experiments were conducted in strict accordance with the recommendations in the Guide for the Care and Use of Laboratory Animals of the National Institutes of Health, USA. The protocol for tick blood-feeding on sheep was approved by the Institutional Animal Care and Use Committee of the University of Southern Mississippi.

## Author Contributions

Conceptualization: Shahid Karim

Methodology: Surendra Raj Sharma, Gary Crispell, Ahmed Mohamed, Cameron Cox, Joshua Lange, Shailesh Chaudhary, Scott Commins, Shahid Karim

Data curation: Surendra Raj Sharma, Gary Crispell, Shailesh Chaudhary, Scott Commins, Shahid Karim

Funding acquisition: Scott P. Commins, Shahid Karim

Investigation: Surendra Sharma, Gary Crispell, Shailesh Chaudhary, Scott Commins, Shahid Karim

Project administration: Shahid Karim Resources: Scott Commins, Shahid Karim Supervision; Shahid Karim

Validation: Surendra Raj Sharma, Scott Commins, Shahid Karim

Writing, original draft: Surendra Raj Sharma, Gary Crispell, Shahid Karim

Writing, review & editing: Surendra Sharma, Scott Commins, Shahid Karim All authors read and approved the manuscript.

## Supplementary information

**Supplementary figure 1.**
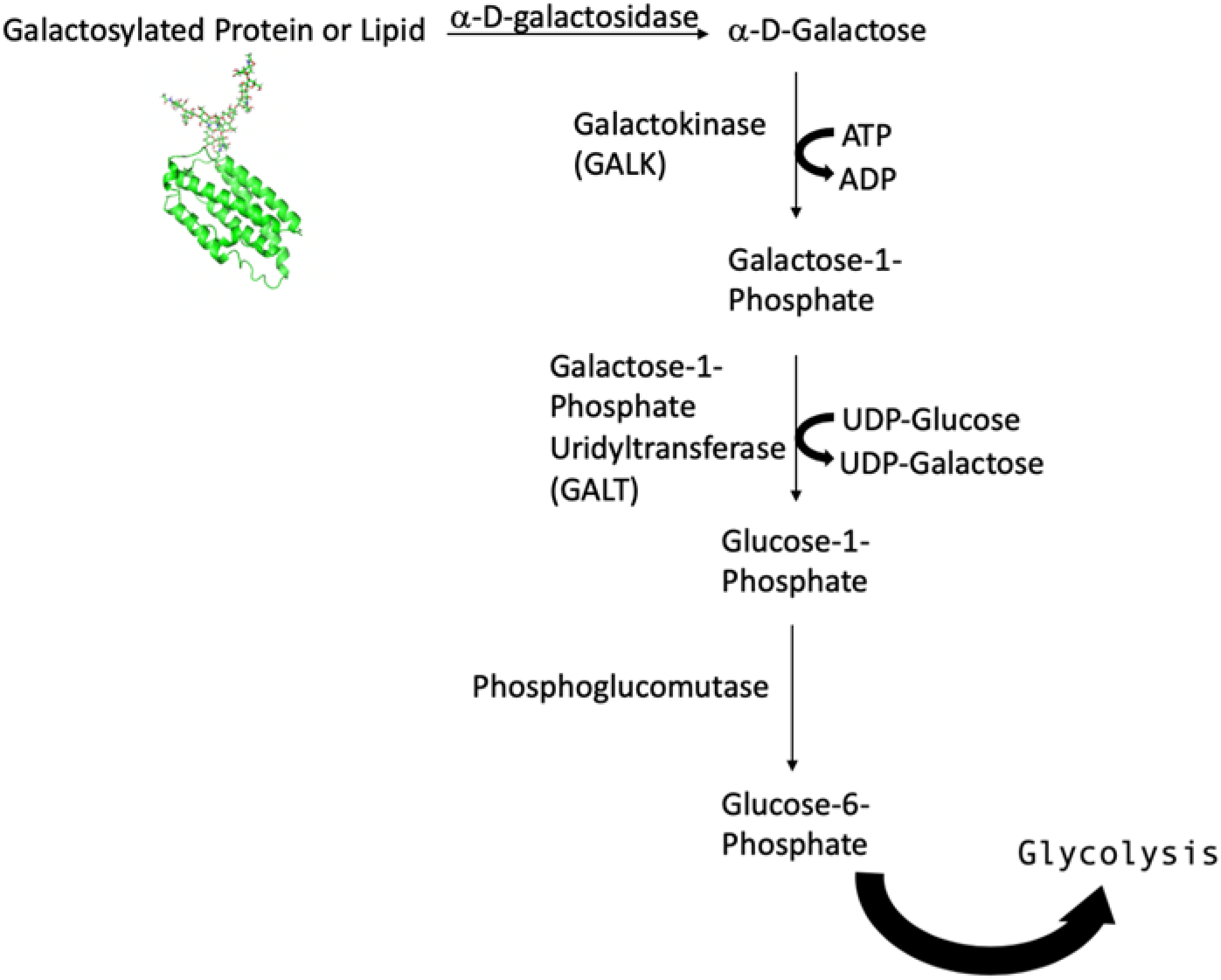
Proposed galactose metabolism pathway of *Amblyomma americanum*.

**Supplementary figure 2:**
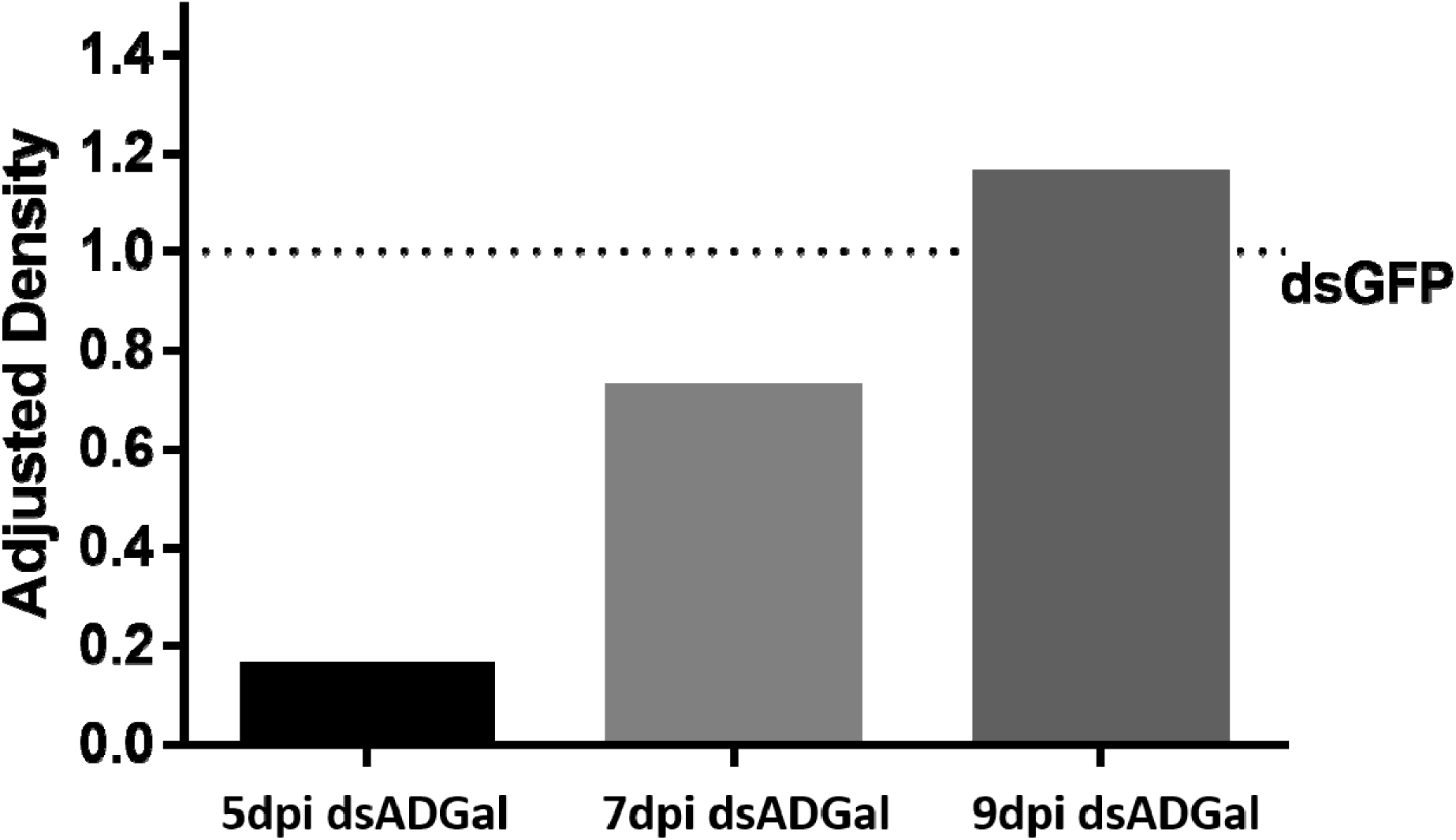
Quantification of relative abundance of α-gal in dsADGal (α-galactosidase) and dsGFP(irrelevant control) injected partially-fed *Am. americanum* salivary glands.

**Supplementary figure 3:**
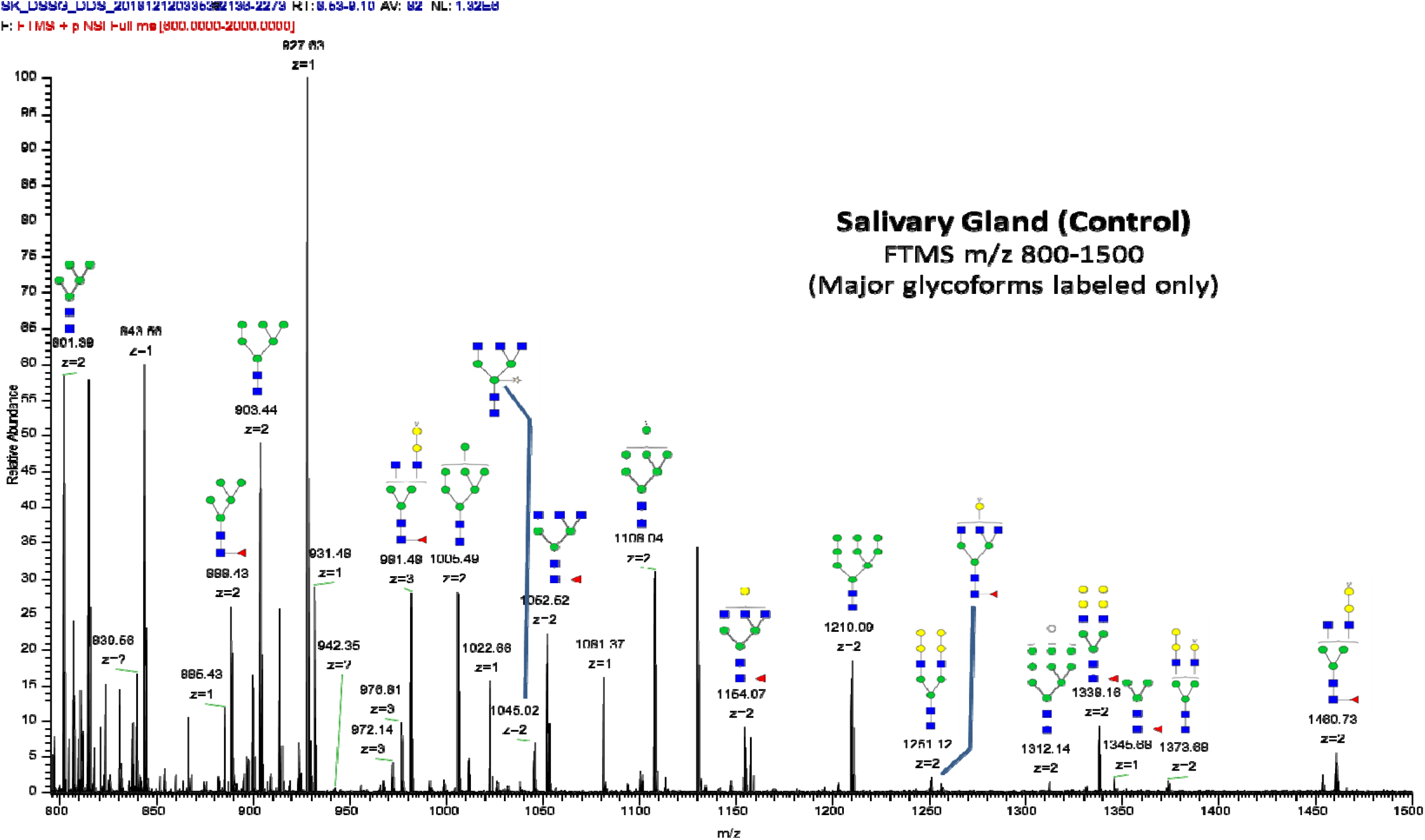
FTMS of N-glycans observed in control Salivary Gland

**Supplementary figure 4:**
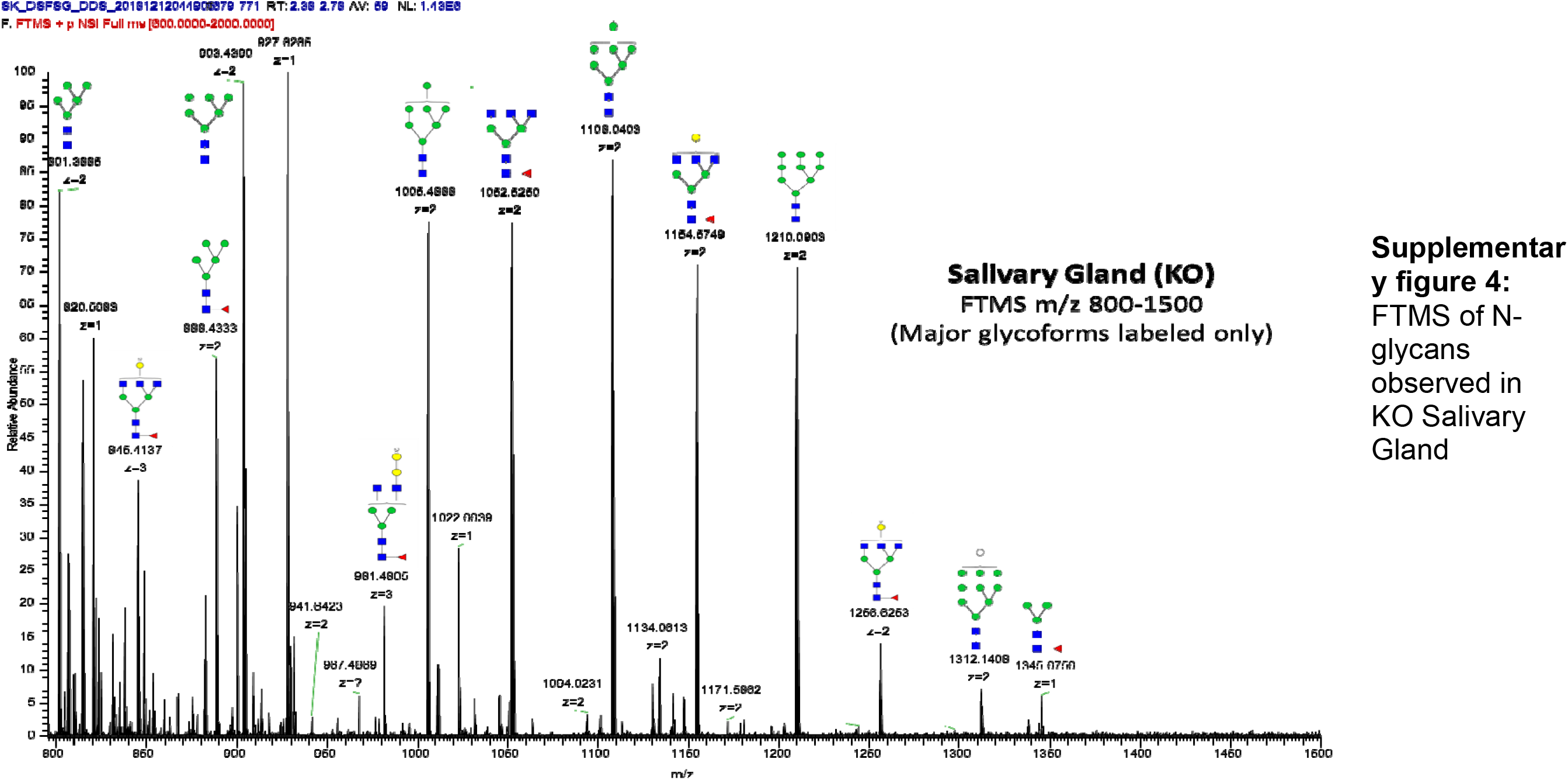
FTMS of N-glycans observed in KO Salivary Gland

**Supplementary figure 5:**
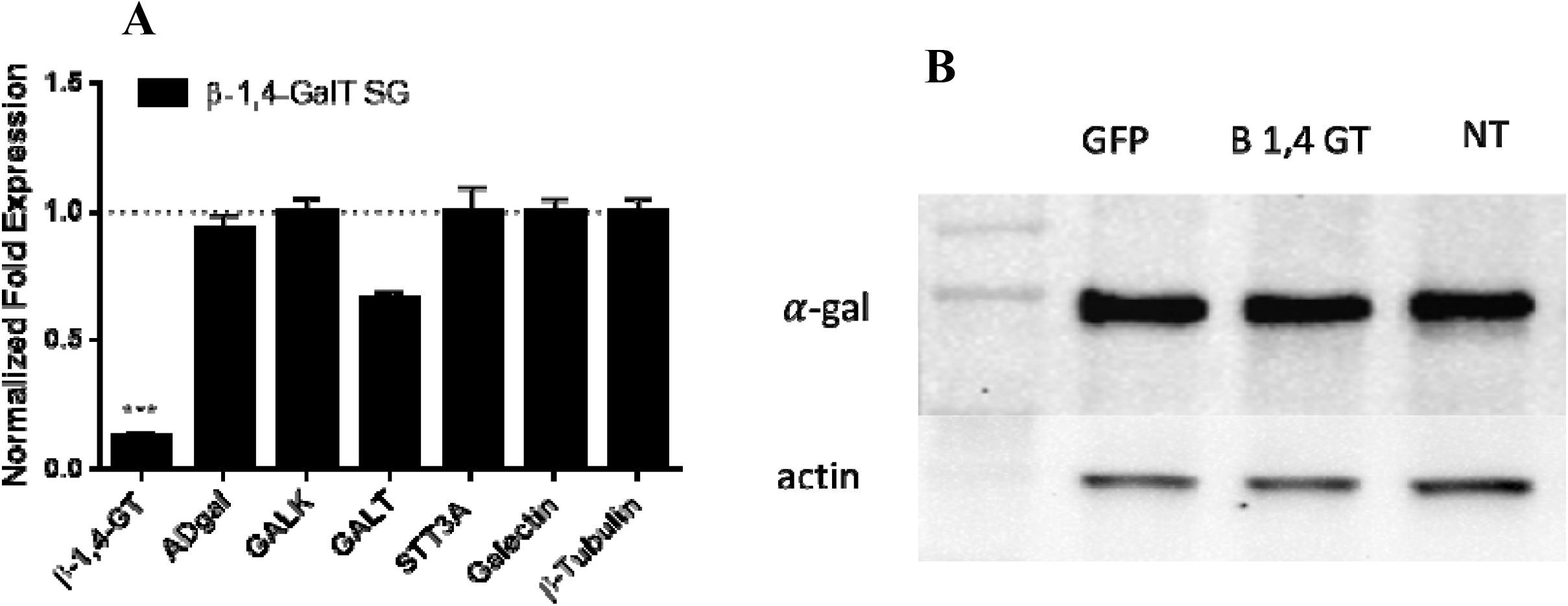
(A) Transcriptional expression of major galactose metabolism, N-glycan synthesis and transport related genes against ß- 1,4 galactosyl transferase silencing in five day partially feed salivary gland(SG) of *Am. americanum*. Actin and Histone was used as house keeping gene and expression was normalized against irrelevant control (GFP dsRNA). Abbreviation: ADgal α-D-galactosidase ,ß- 1,4 GT: ß- 1,4 galactosyl transferase; GALK: Galacto kinase; GALT galactosyl transferase; STT3A: Dolichyl-diphosphooligosaccharide--protein glycosyltransferase(***P<0.001, student t test) (B) Detection of α-gal in ds ß-1,4GT, dsGFP and non treated injected partially-fed tick salivary glands

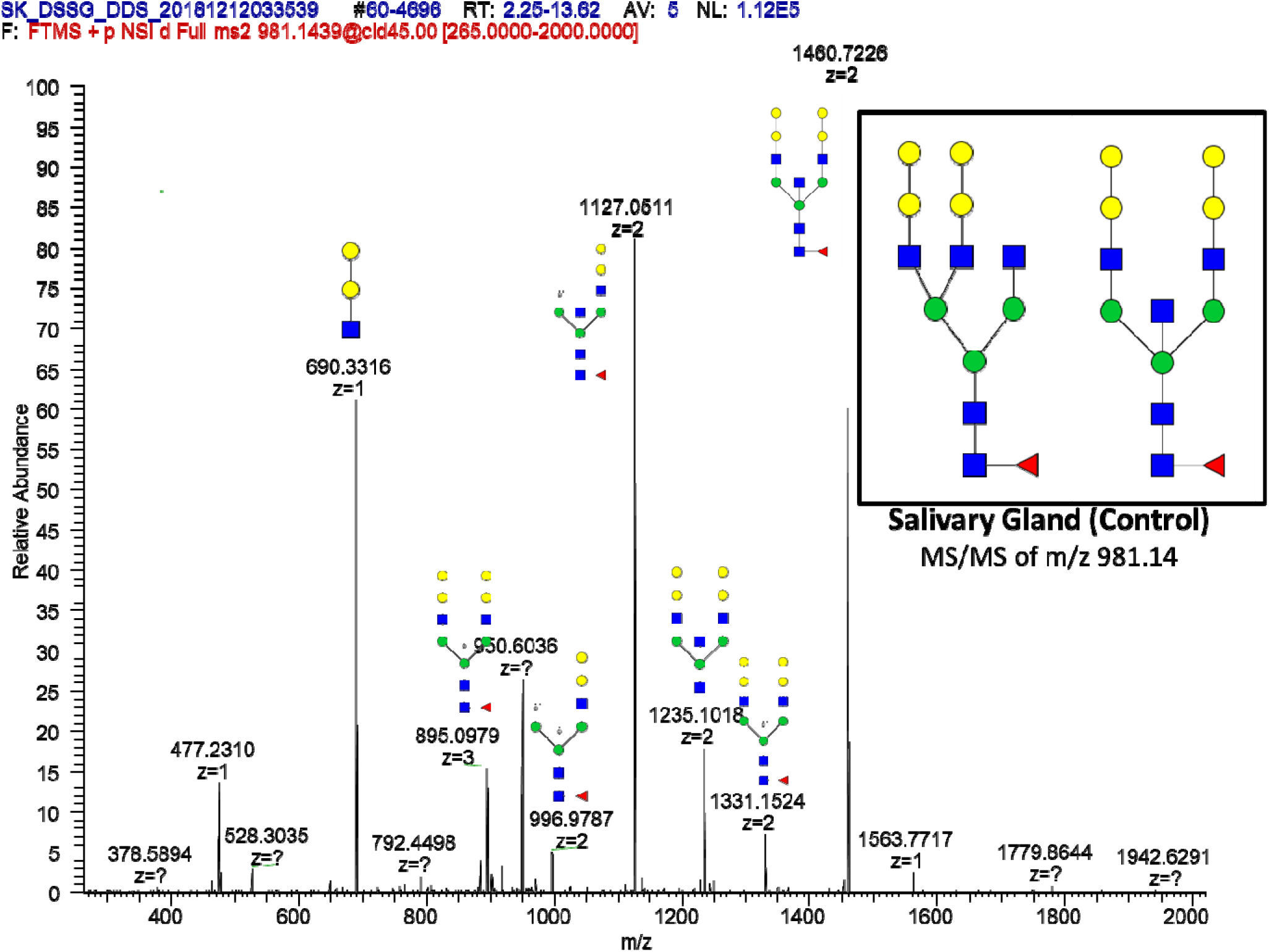
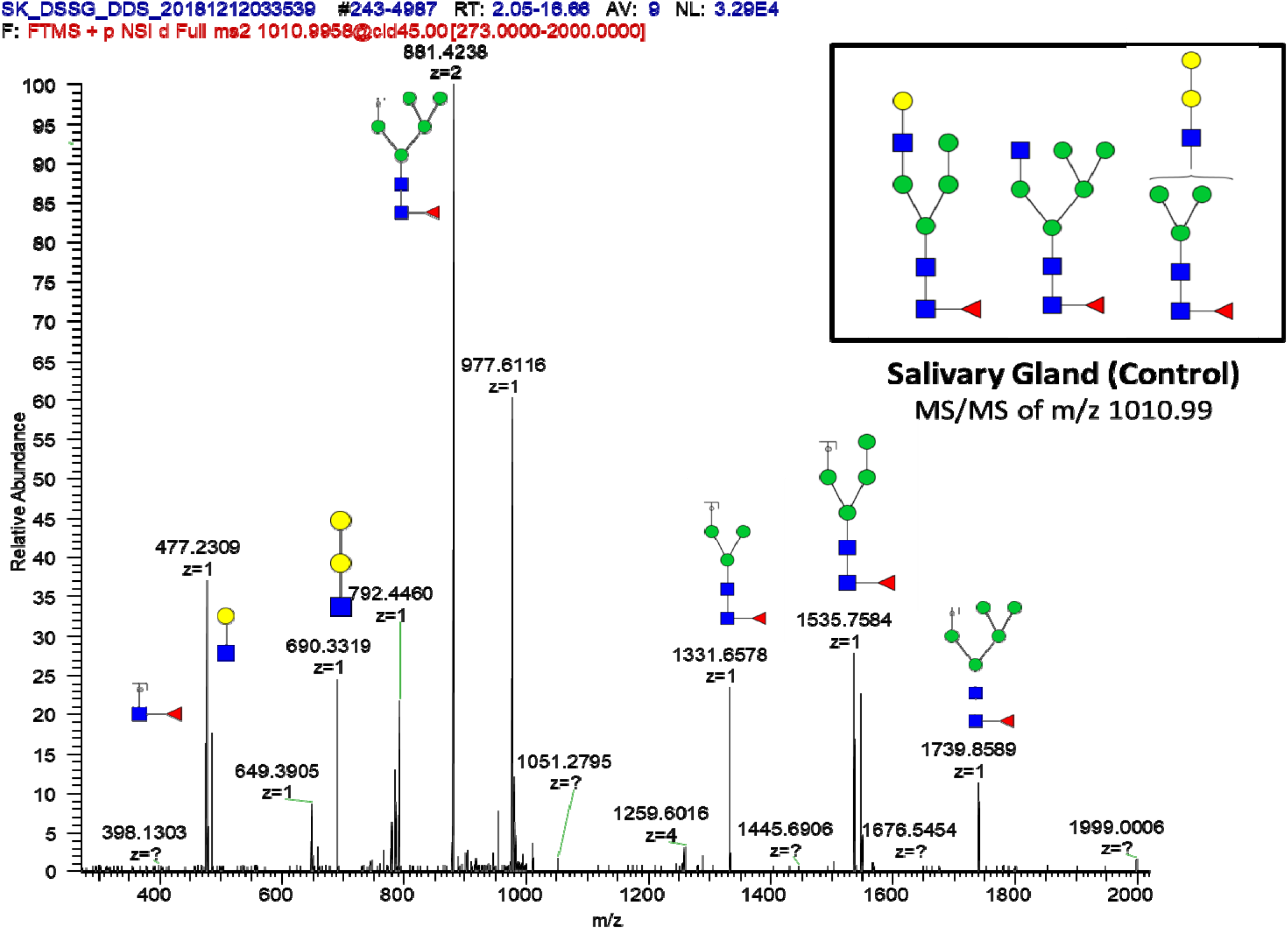
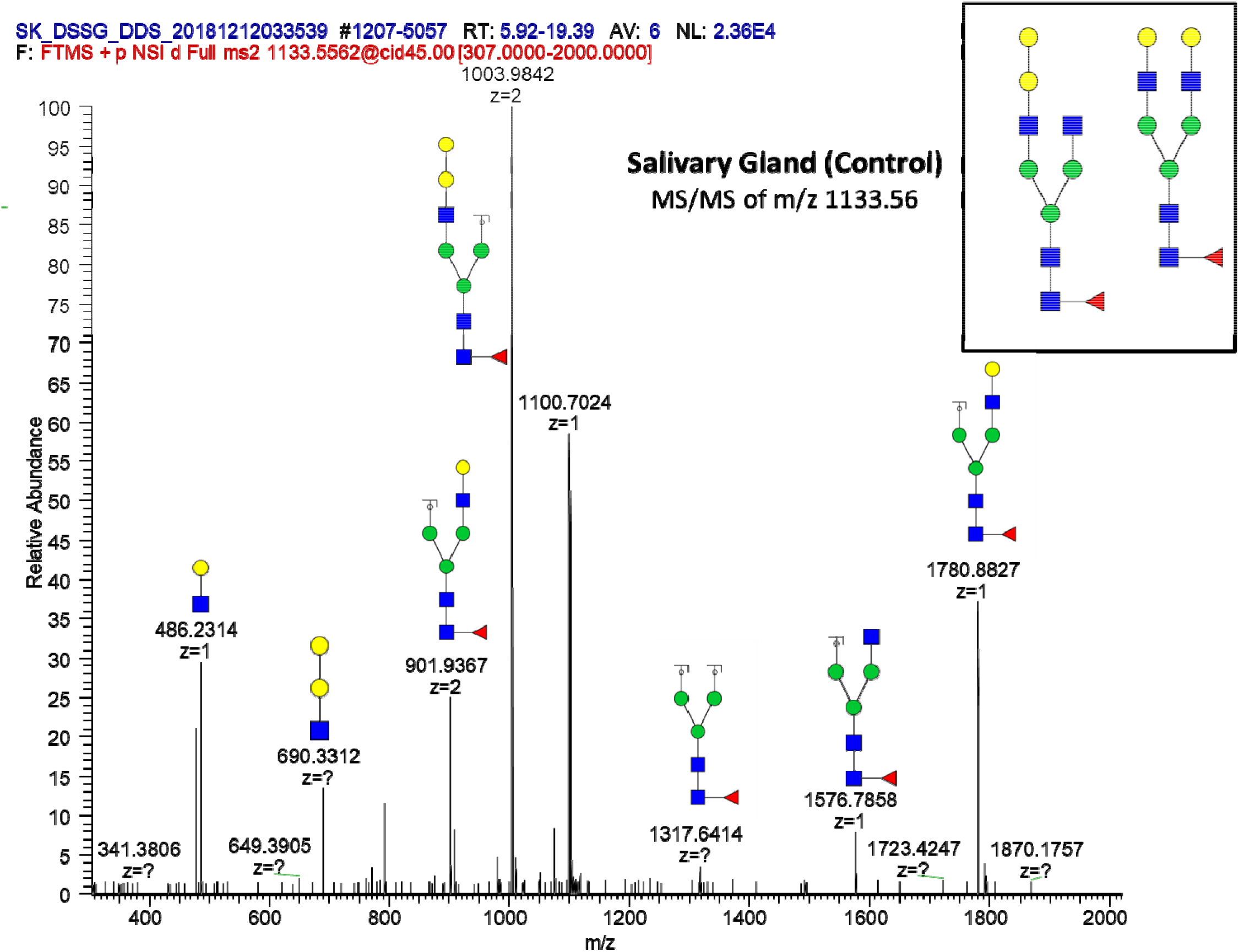
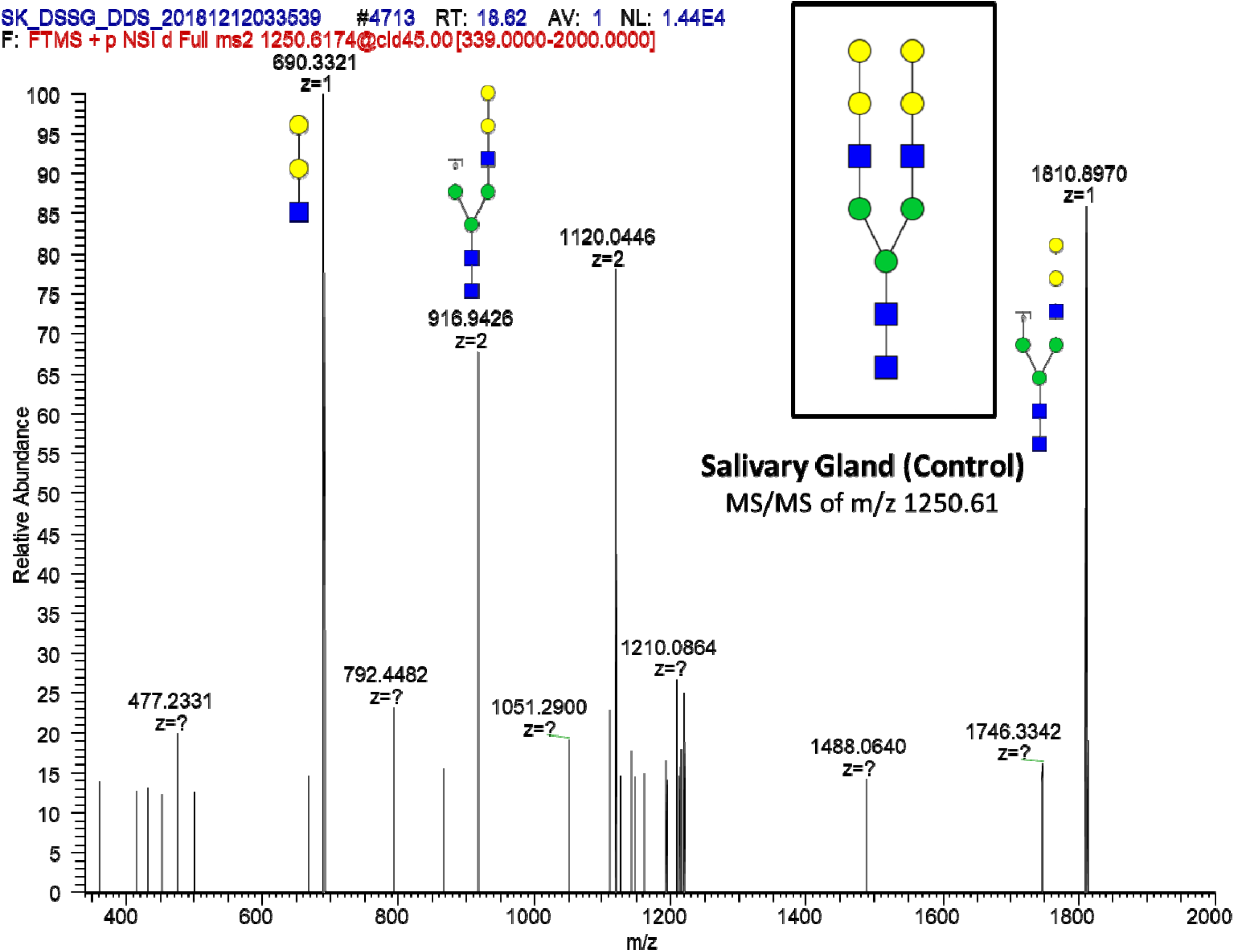
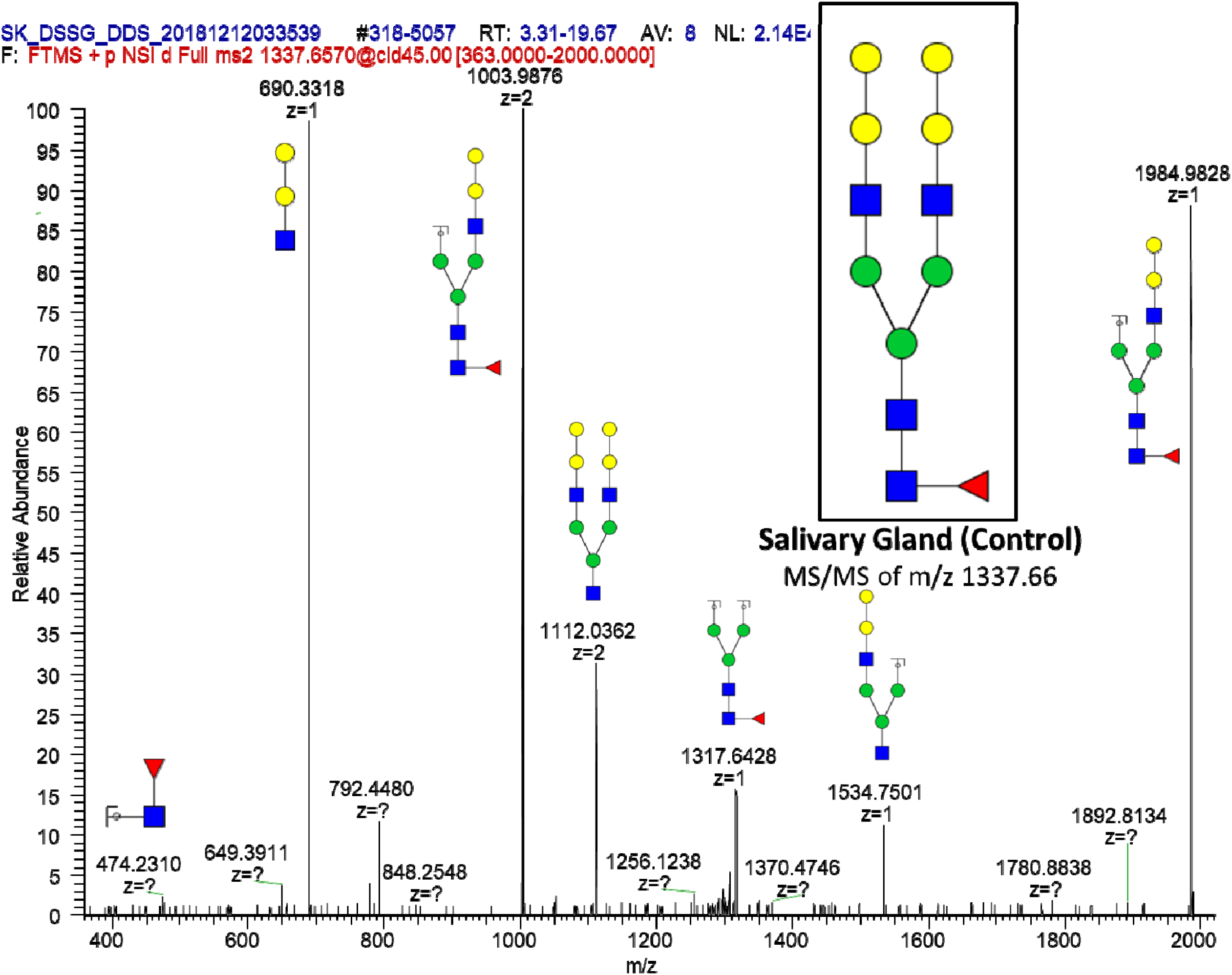
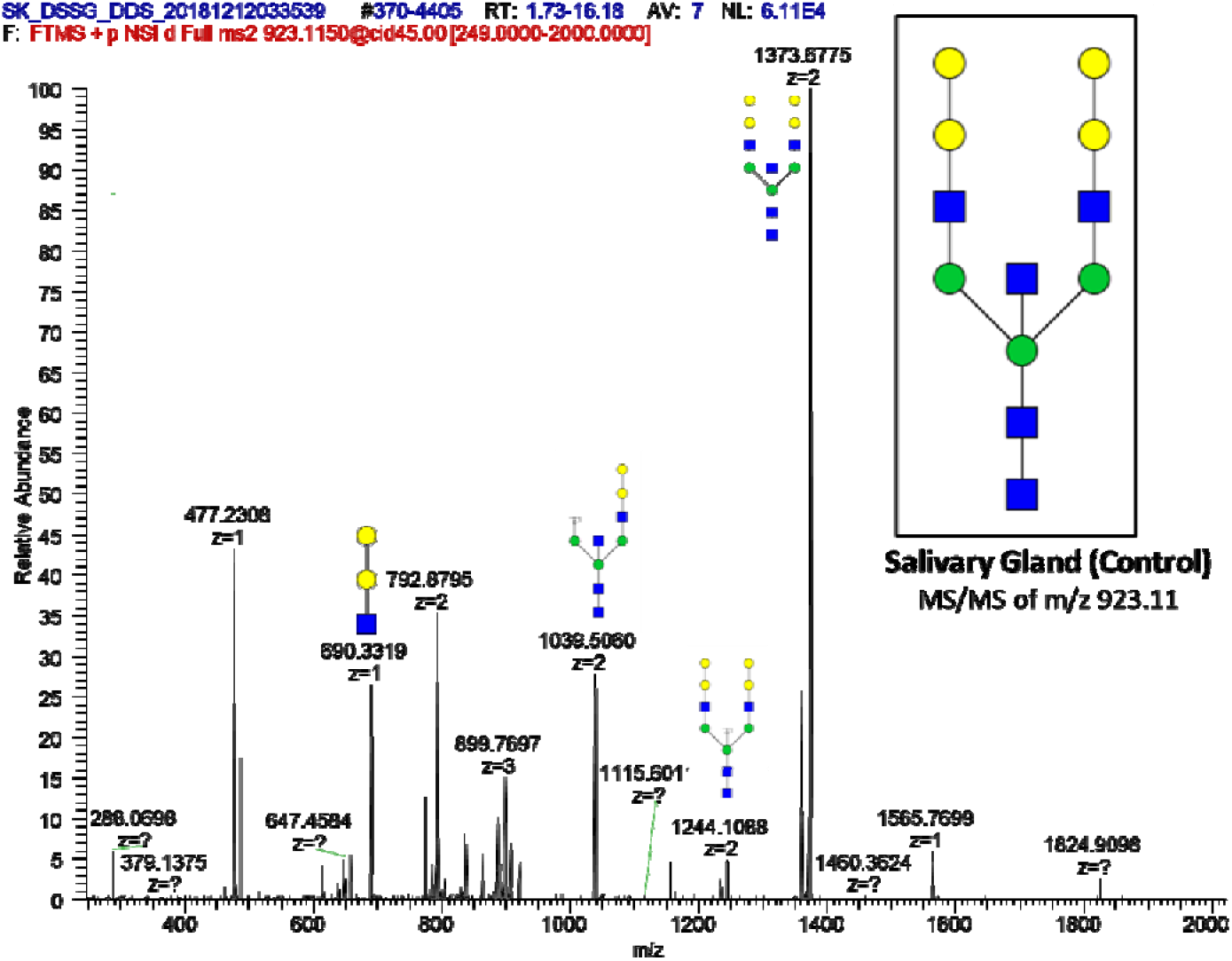
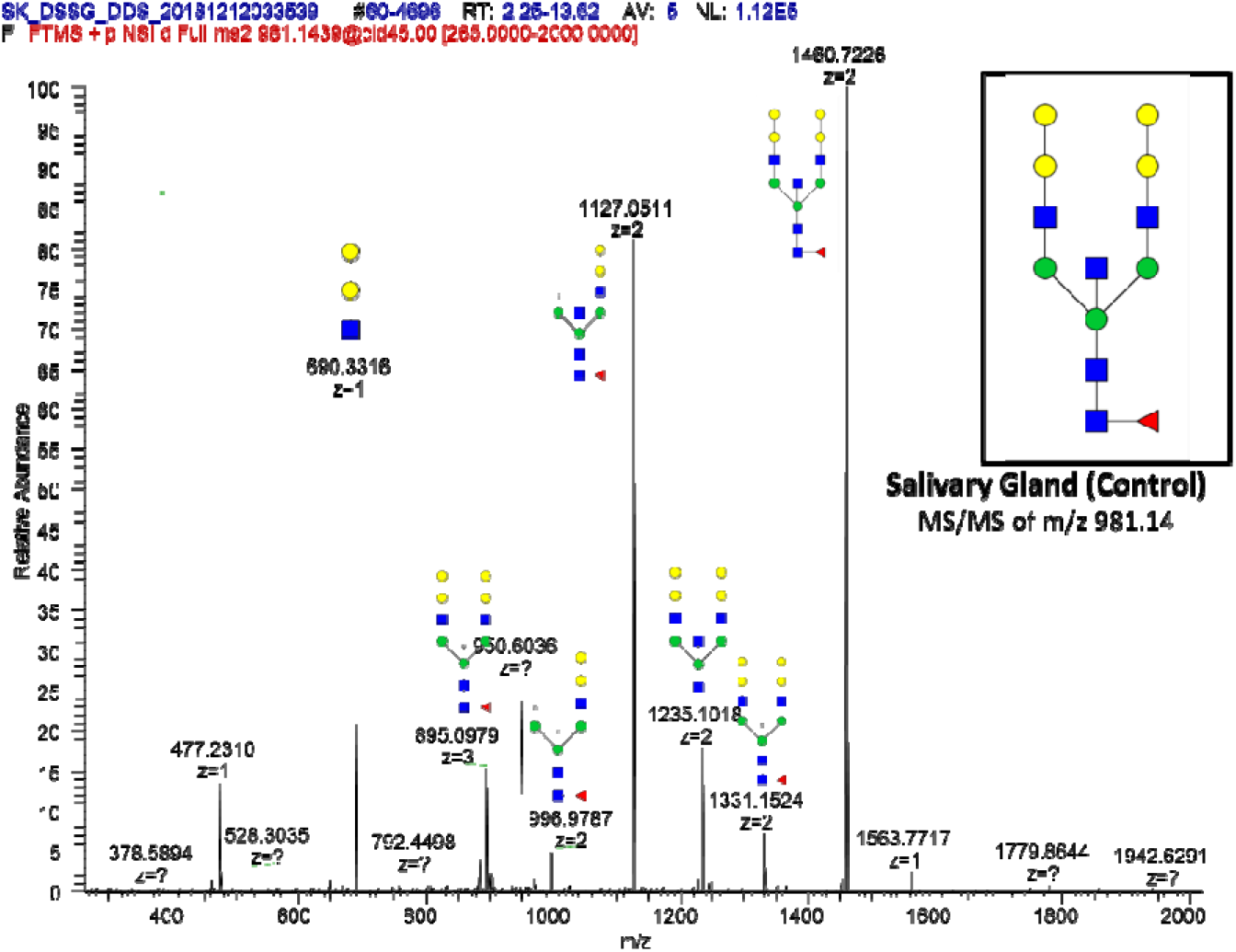
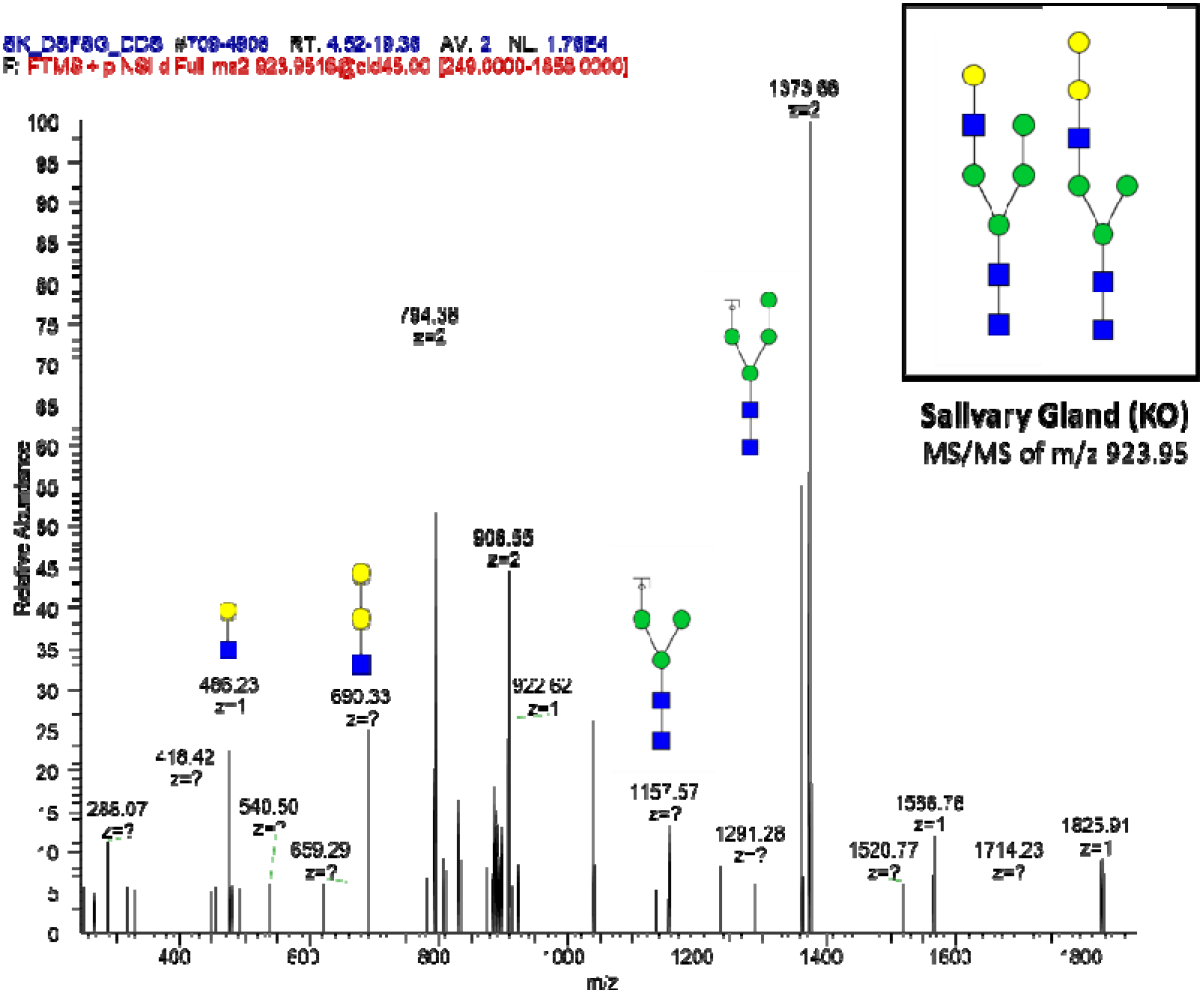
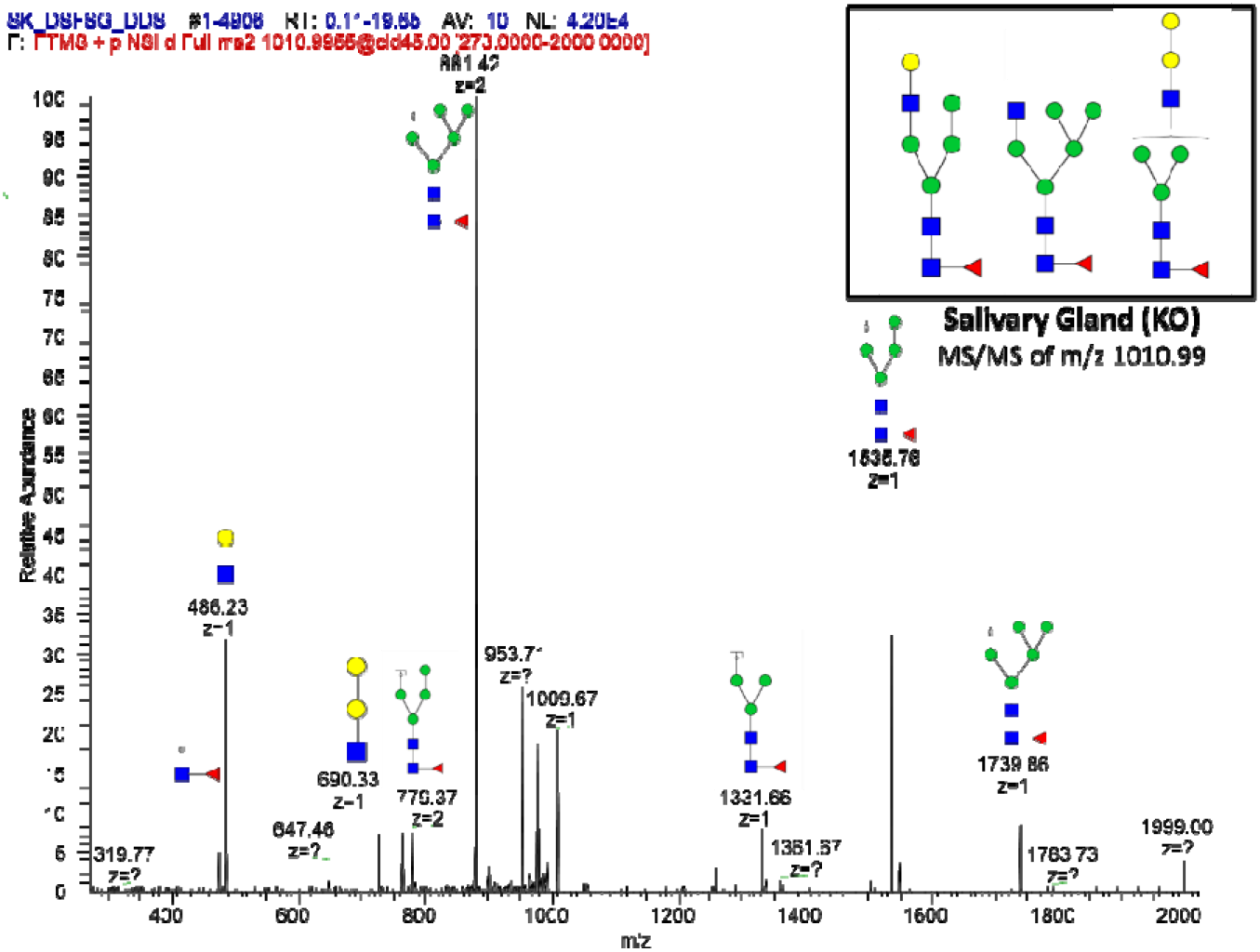
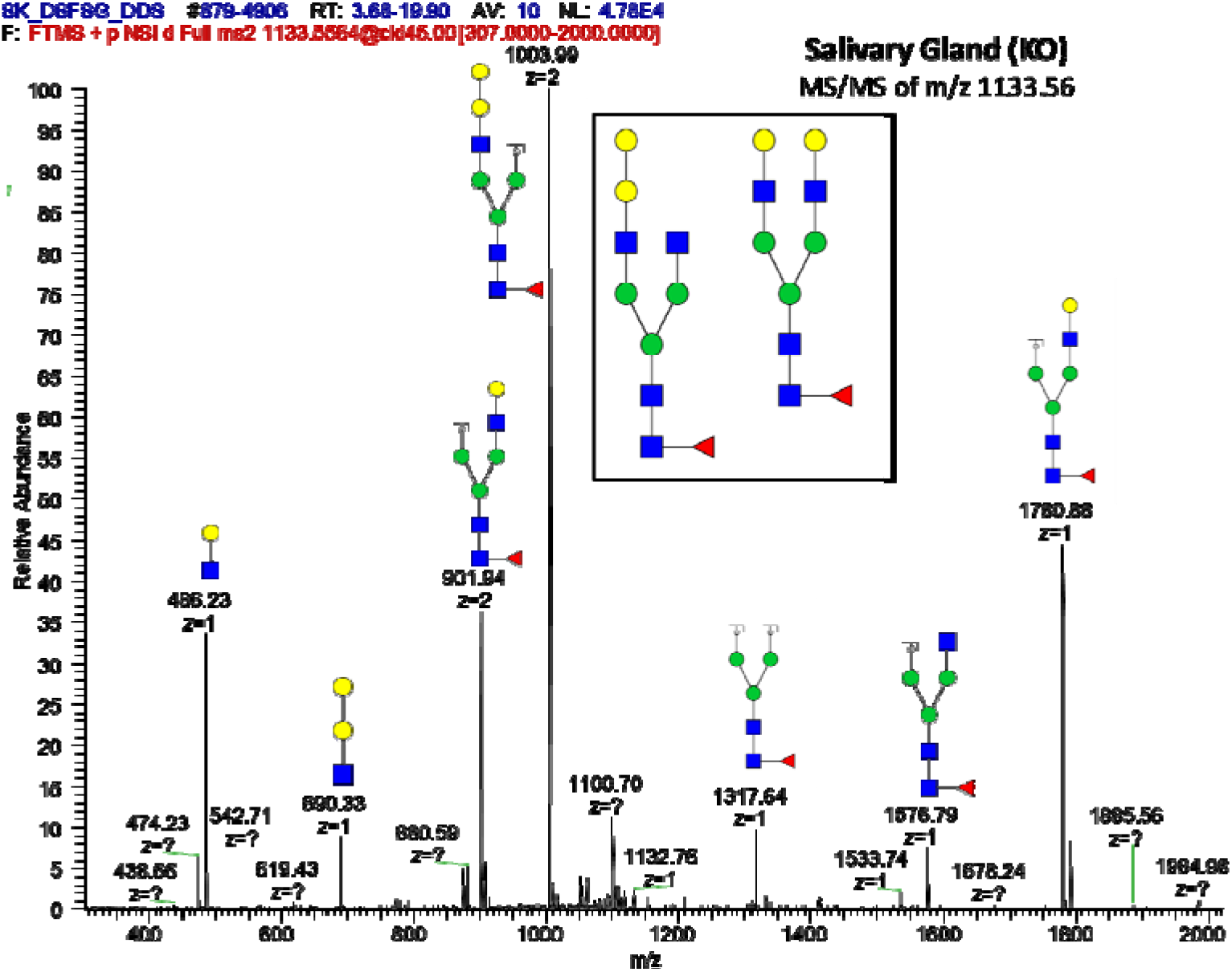
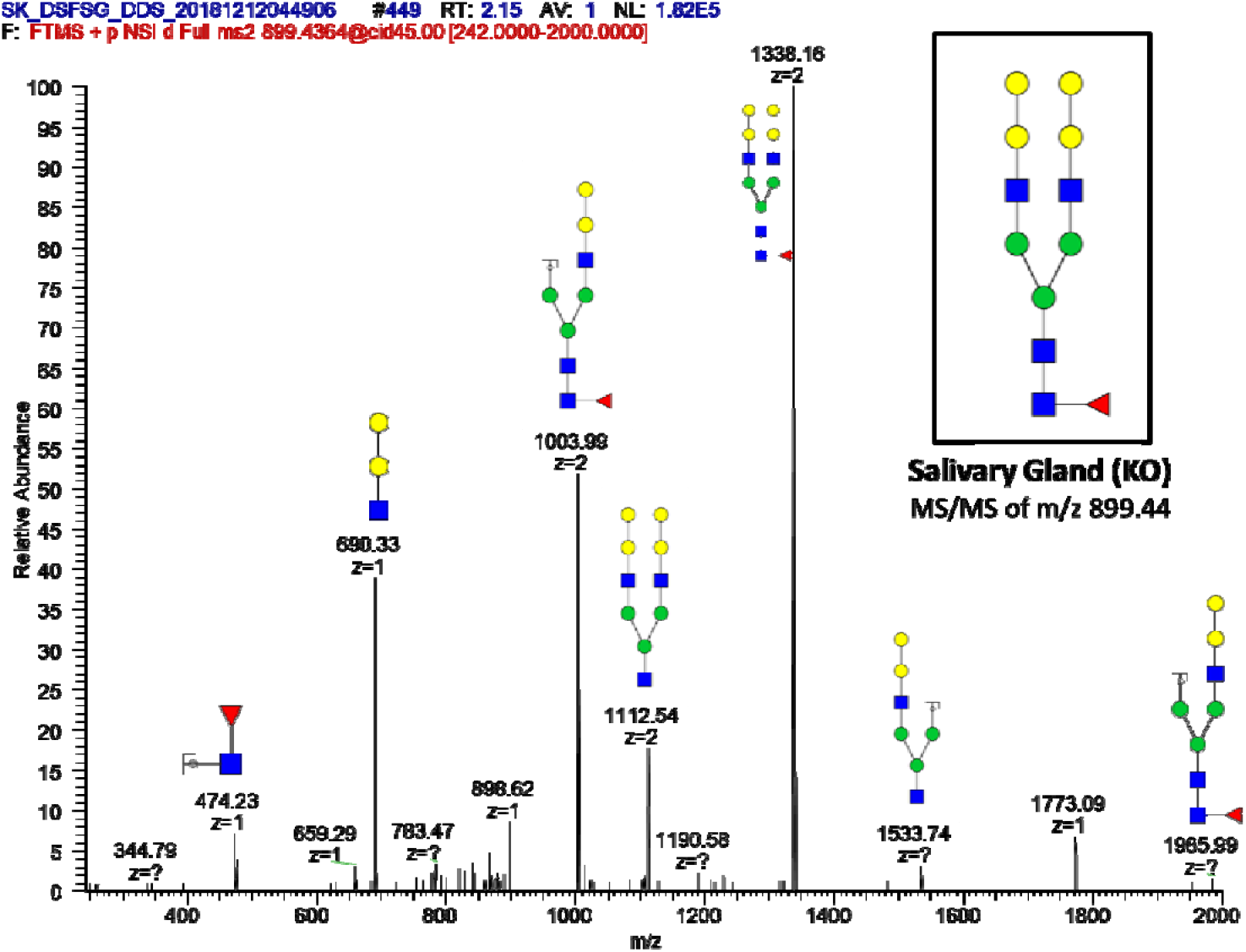
Miscellaneous supplementary figures: FTMS of all N-glycans observed in KO Salivary and control Gland

**Supplementary Table 1:**
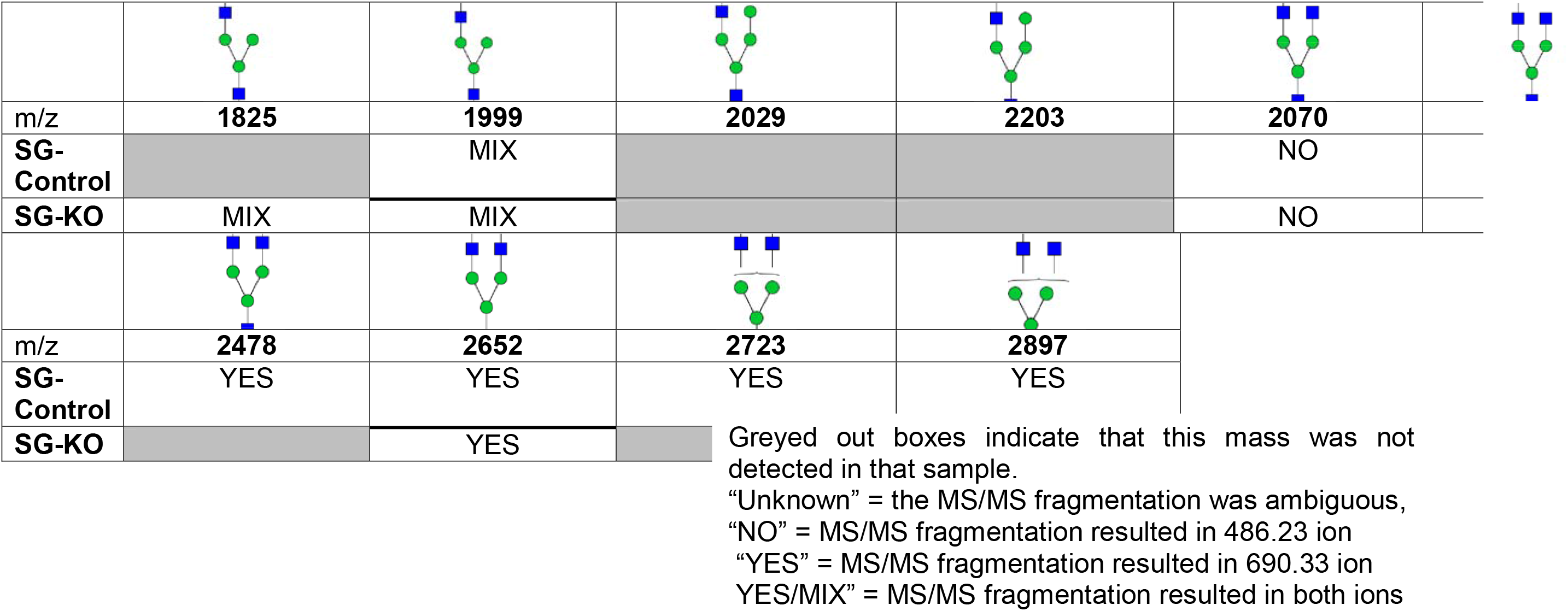
Summary of potential α-Gal Glycoforms Determined by MS/MS Analysis

|  |  |  |  |  |  |
| --- | --- | --- | --- | --- | --- |
| m/z | **1825** | **1999** | **2029** | **2203** | **2070** |
| SG-Control |  | MIX |  |  | NO |
| SG-KO | MIX | MIX |  |  | NO |
| m/z | **2478** | **2652** | **2723** | **2897** |  |
| SG-Control | YES | YES | YES | YES |  |
| SG-KO |  | YES |  |  |  |

Greyed out boxes indicate that this mass was not detected in that sample.
“Unknown” = the MS/MS fragmentation was ambiguous,
“NO” = MS/MS fragmentation resulted in 486.23 ion
“YES” = MS/MS fragmentation resulted in 690.33 ion
YES/MIX” = MS/MS fragmentation resulted in both ions

**Supplementary table 2:**
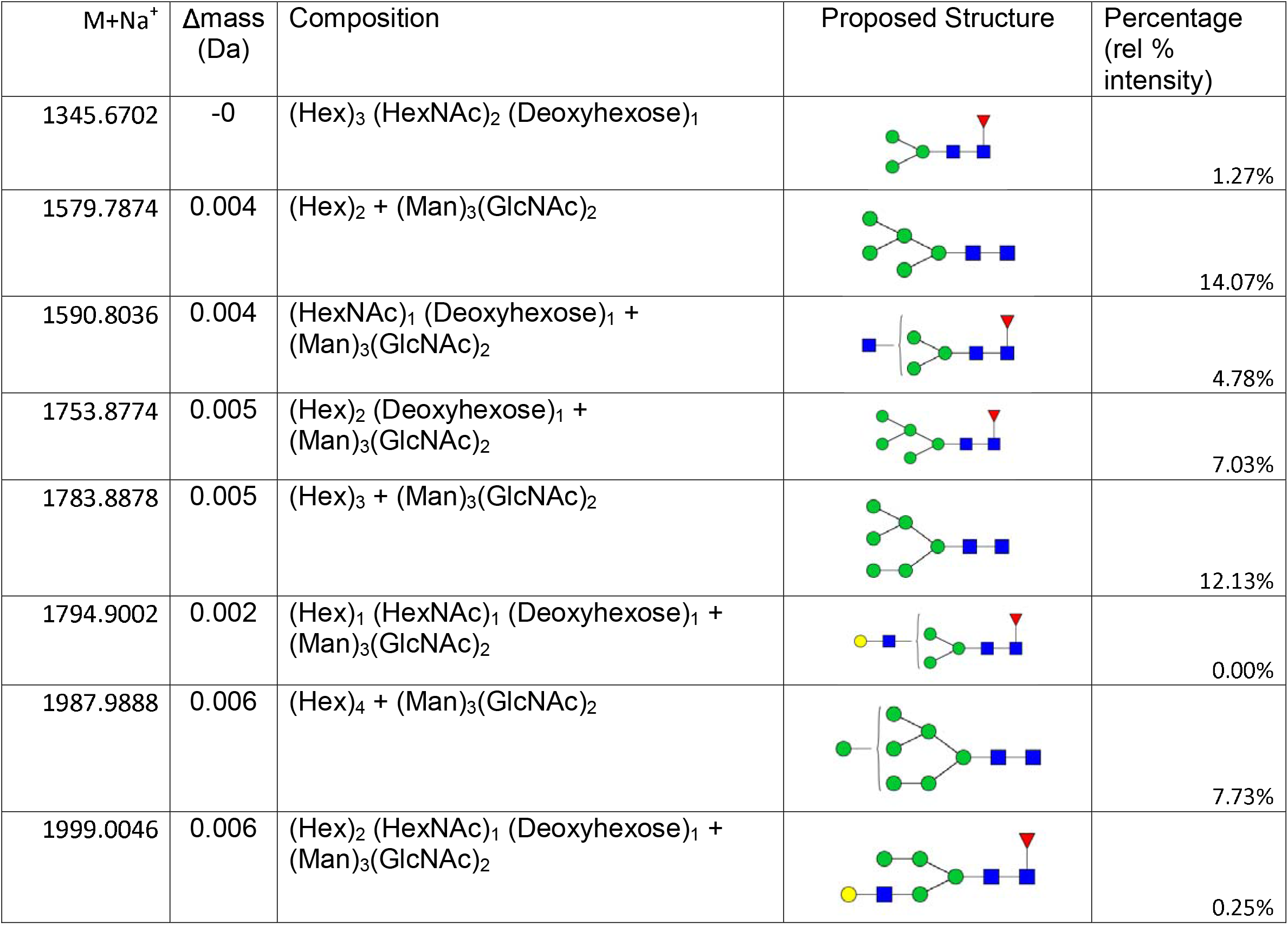
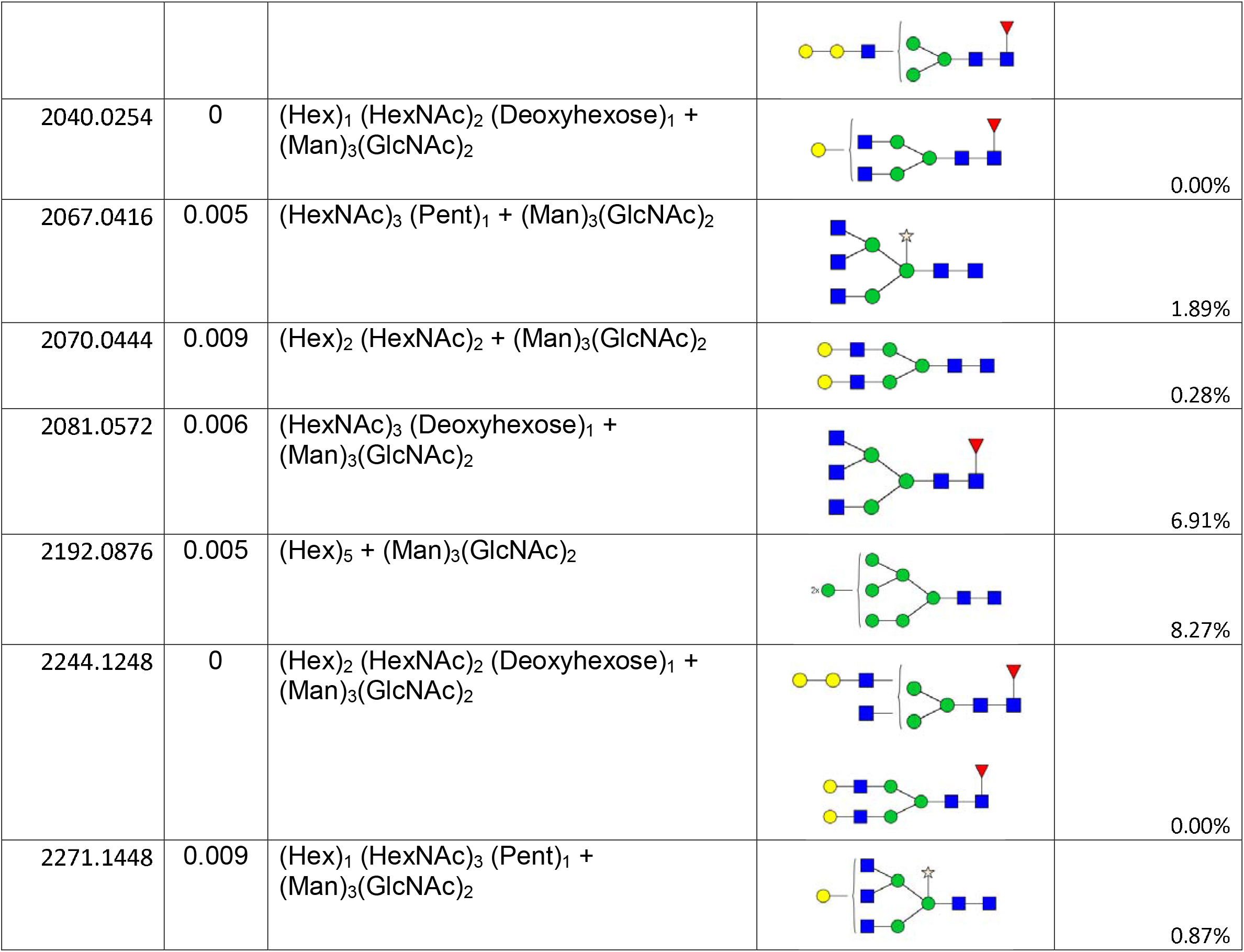
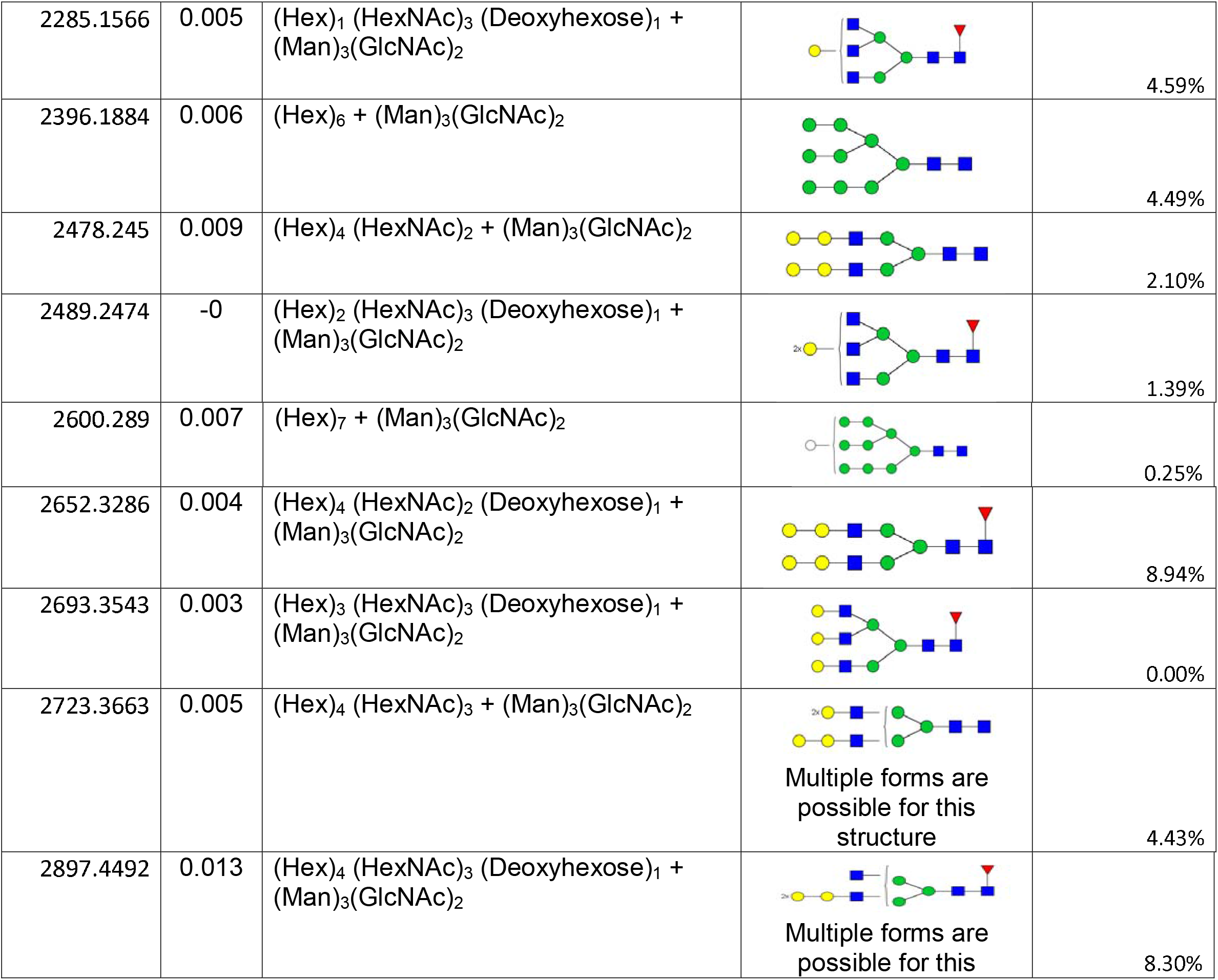
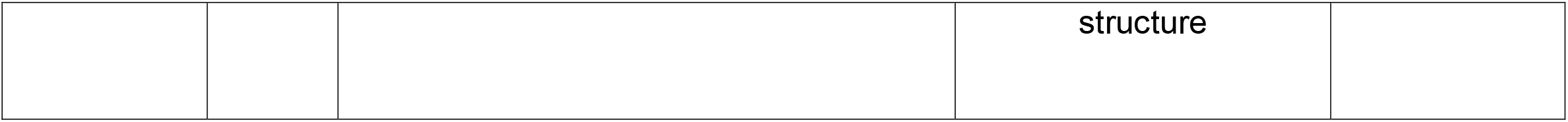
FTMS of N-Glycans observed in control salivary glands.

| M+Na<sup>+</sup> | Δmass (Da) | Composition | Proposed Structure | Percentage (rel % intensity) |
| --- | --- | --- | --- | --- |
| 1345.6702 | -0 | (Hex)<sub>3</sub> (HexNAc)<sub>2</sub> (Deoxyhexose)<sub>1</sub> |  | 1.27% |
| 1579.7874 | 0.004 | (Hex)<sub>2</sub> + (Man)<sub>3</sub>(GlcNAc)<sub>2</sub> |  | 14.07% |
| 1590.8036 | 0.004 | (HexNAc)<sub>1</sub> (Deoxyhexose)<sub>1</sub> + (Man)<sub>3</sub>(GlcNAc)<sub>2</sub> |  | 4.78% |
| 1753.8774 | 0.005 | (Hex)<sub>2</sub> (Deoxyhexose)<sub>1</sub> + (Man)<sub>3</sub>(GlcNAc)<sub>2</sub> |  | 7.03% |
| 1783.8878 | 0.005 | (Hex)<sub>3</sub> + (Man)<sub>3</sub>(GlcNAc)<sub>2</sub> |  | 12.13% |
| 1794.9002 | 0.002 | (Hex)<sub>1</sub> (HexNAc)<sub>1</sub> (Deoxyhexose)<sub>1</sub> + (Man)<sub>3</sub>(GlcNAc)<sub>2</sub> |  | 0.00% |
| 1987.9888 | 0.006 | (Hex)<sub>4</sub> + (Man)<sub>3</sub>(GlcNAc)<sub>2</sub> |  | 7.73% |
| 1999.0046 | 0.006 | (Hex)<sub>2</sub> (HexNAc)<sub>1</sub> (Deoxyhexose)<sub>1</sub> + (Man)<sub>3</sub>(GlcNAc)<sub>2</sub> |  | 0.25% |
| 2040.0254 | 0 | (Hex)<sub>1</sub> (HexNAc)<sub>2</sub> (Deoxyhexose)<sub>1</sub> + (Man)<sub>3</sub>(GlcNAc)<sub>2</sub> |  | 0.00% |
| 2067.0416 | 0.005 | (HexNAc)<sub>3</sub> (Pent)<sub>1</sub> + (Man)<sub>3</sub>(GlcNAc)<sub>2</sub> |  | 1.89% |
| 2070.0444 | 0.009 | (Hex)<sub>2</sub> (HexNAc)<sub>2</sub> + (Man)<sub>3</sub>(GlcNAc)<sub>2</sub> |  | 0.28% |
| 2081.0572 | 0.006 | (HexNAc)<sub>3</sub> (Deoxyhexose)<sub>1</sub> + (Man)<sub>3</sub>(GlcNAc)<sub>2</sub> |  | 6.91% |
| 2192.0876 | 0.005 | (Hex)<sub>5</sub> + (Man)<sub>3</sub>(GlcNAc)<sub>2</sub> |  | 8.27% |
| 2244.1248 | 0 | (Hex)<sub>2</sub> (HexNAc)<sub>2</sub> (Deoxyhexose)<sub>1</sub> + (Man)<sub>3</sub>(GlcNAc)<sub>2</sub> |  | 0.00% |
| 2271.1448 | 0.009 | (Hex)<sub>1</sub> (HexNAc)<sub>3</sub> (Pent)<sub>1</sub> + (Man)<sub>3</sub>(GlcNAc)<sub>2</sub> |  | 0.87% |
| 2285.1566 | 0.005 | (Hex)<sub>1</sub> (HexNAc)<sub>3</sub> (Deoxyhexose)<sub>1</sub> + (Man)<sub>3</sub>(GlcNAc)<sub>2</sub> |  | 4.59% |
| 2396.1884 | 0.006 | (Hex)<sub>6</sub> + (Man)<sub>3</sub>(GlcNAc)<sub>2</sub> |  | 4.49% |
| 2478.245 | 0.009 | (Hex)<sub>4</sub> (HexNAc)<sub>2</sub> + (Man)<sub>3</sub>(GlcNAc)<sub>2</sub> |  | 2.10% |
| 2489.2474 | -0 | (Hex)<sub>2</sub> (HexNAc)<sub>3</sub> (Deoxyhexose)<sub>1</sub> + (Man)<sub>3</sub>(GlcNAc)<sub>2</sub> |  | 1.39% |
| 2600.289 | 0.007 | (Hex)<sub>7</sub> + (Man)<sub>3</sub>(GlcNAc)<sub>2</sub> |  | 0.25% |
| 2652.3286 | 0.004 | (Hex)<sub>4</sub> (HexNAc)<sub>2</sub> (Deoxyhexose)<sub>1</sub> + (Man)<sub>3</sub>(GlcNAc)<sub>2</sub> |  | 8.94% |
| 2693.3543 | 0.003 | (Hex)<sub>3</sub> (HexNAc)<sub>3</sub> (Deoxyhexose)<sub>1</sub> + (Man)<sub>3</sub>(GlcNAc)<sub>2</sub> |  | 0.00% |
| 2723.3663 | 0.005 | (Hex)<sub>4</sub> (HexNAc)<sub>3</sub> + (Man)<sub>3</sub>(GlcNAc)<sub>2</sub> | Multiple forms are possible for this structure | 4.43% |
| 2897.4492 | 0.013 | (Hex)<sub>4</sub> (HexNAc)<sub>3</sub> (Deoxyhexose)<sub>1</sub> + (Man)<sub>3</sub>(GlcNAc)<sub>2</sub> | Multiple forms are possible for this structure | 8.30% |

**Supplementary Table 3:**
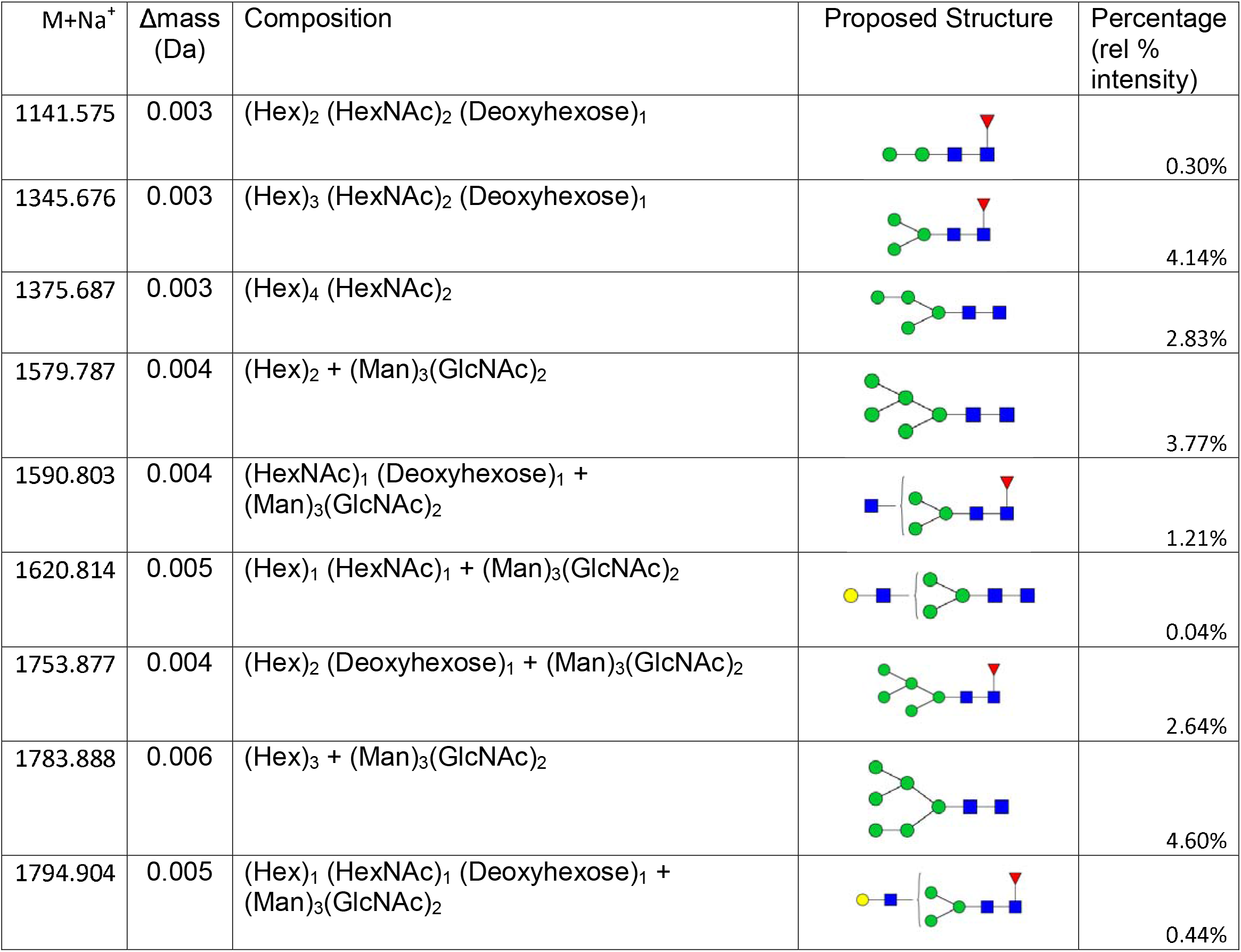
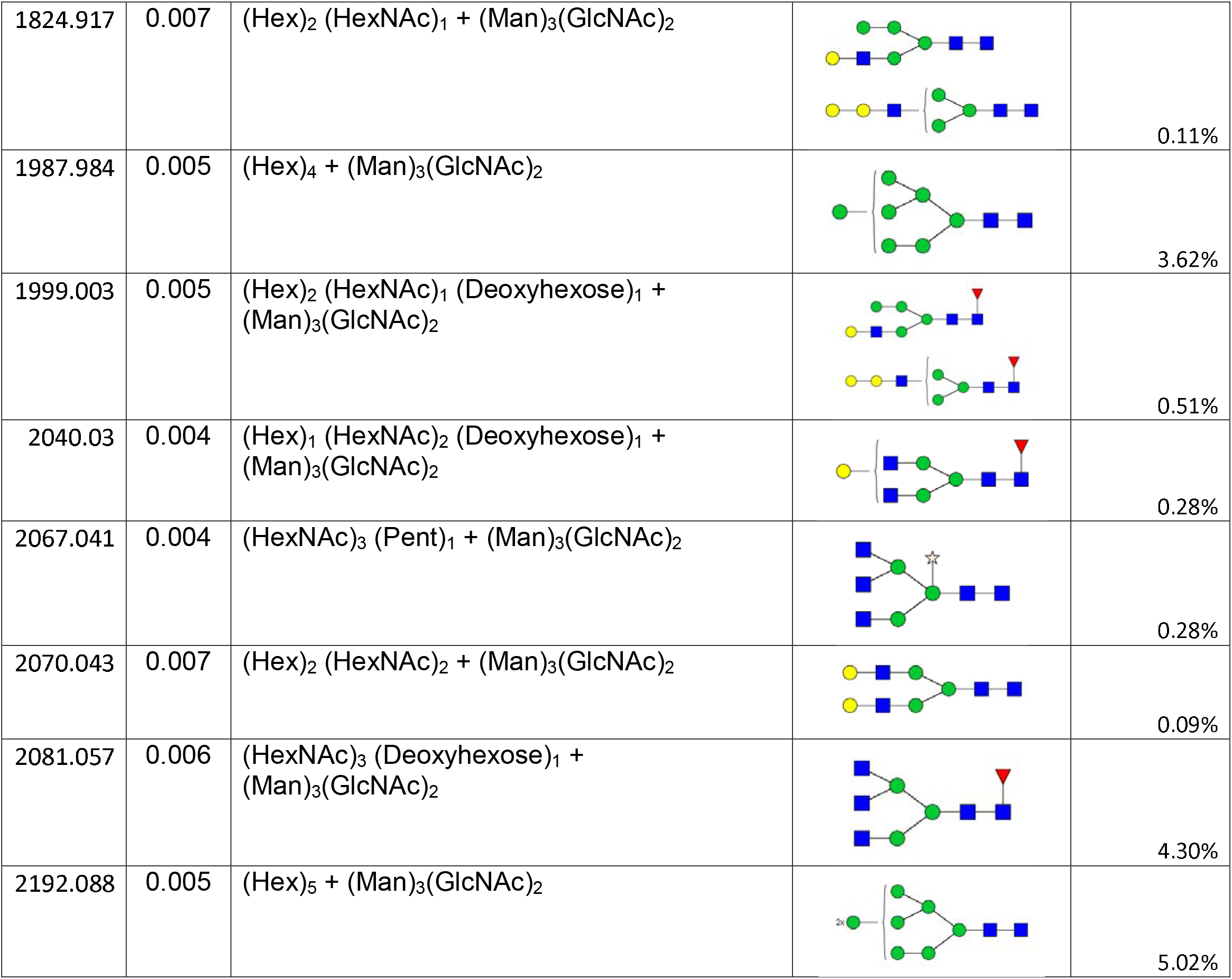
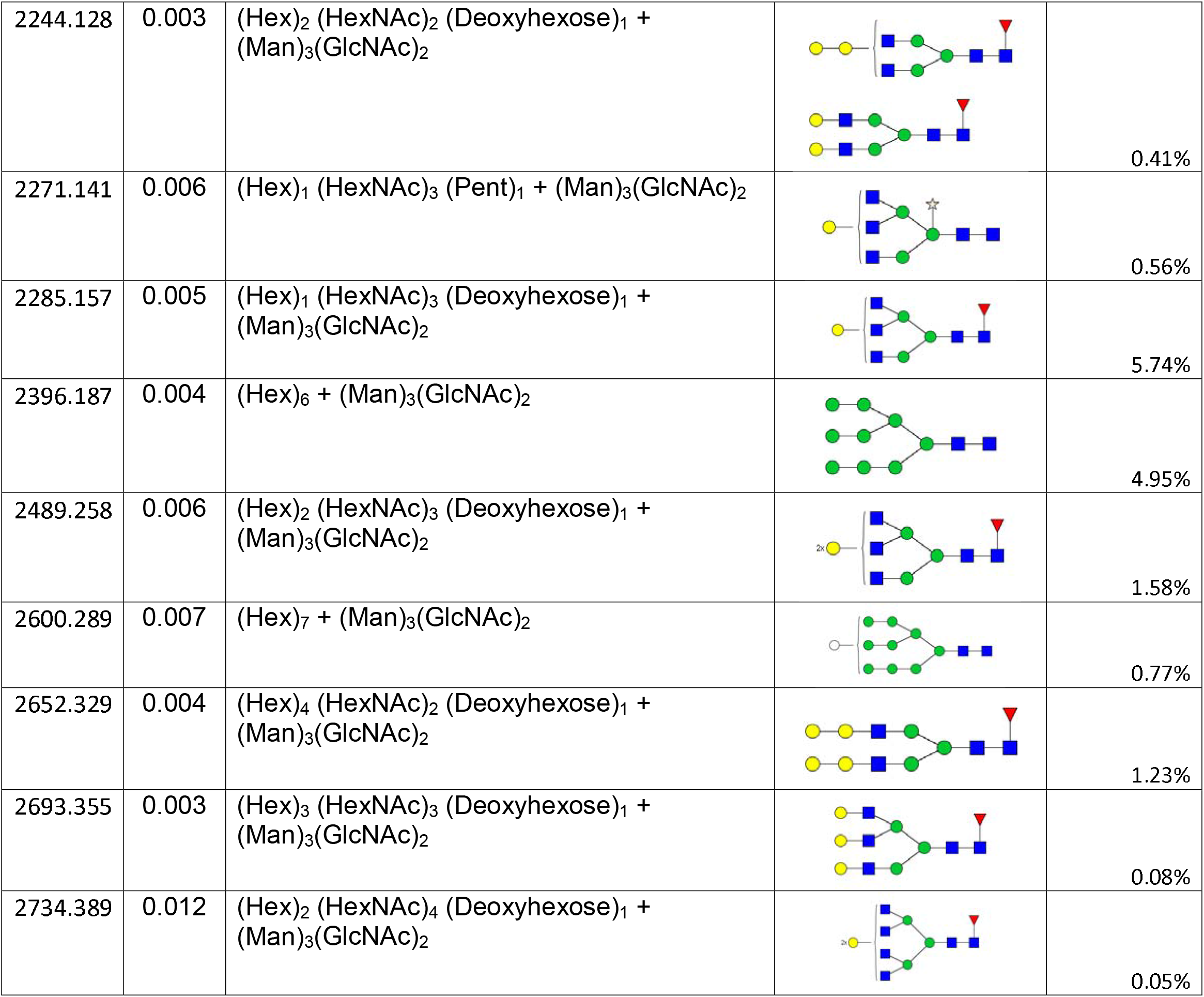
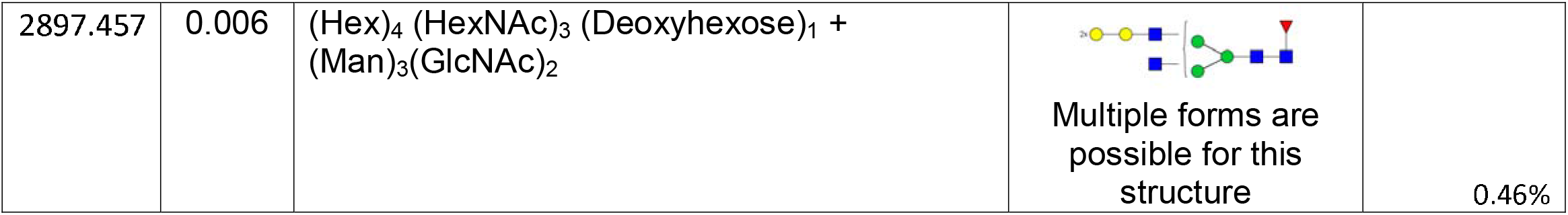
FTMS of N-glycans observed in α-D-galactosidase silenced salivary glands

| M+Na<sup>+</sup> | Δmass (Da) | Composition | Proposed Structure | Percentage (rel % intensity) |
| --- | --- | --- | --- | --- |
| 1141.575 | 0.003 | (Hex)<sub>2</sub> (HexNAc)<sub>2</sub> (Deoxyhexose)<sub>1</sub> |  | 0.30% |
| 1345.676 | 0.003 | (Hex)<sub>3</sub> (HexNAc)<sub>2</sub> (Deoxyhexose)<sub>1</sub> |  | 4.14% |
| 1375.687 | 0.003 | (Hex)<sub>4</sub> (HexNAc)<sub>2</sub> |  | 2.83% |
| 1579.787 | 0.004 | (Hex)<sub>2</sub> + (Man)<sub>3</sub>(GlcNAc)<sub>2</sub> |  | 3.77% |
| 1590.803 | 0.004 | (HexNAc)<sub>1</sub> (Deoxyhexose)<sub>1</sub> + (Man)<sub>3</sub>(GlcNAc)<sub>2</sub> |  | 1.21% |
| 1620.814 | 0.005 | (Hex)<sub>1</sub> (HexNAc)<sub>1</sub> + (Man)<sub>3</sub>(GlcNAc)<sub>2</sub> |  | 0.04% |
| 1753.877 | 0.004 | (Hex)<sub>2</sub> (Deoxyhexose)<sub>1</sub> + (Man)<sub>3</sub>(GlcNAc)<sub>2</sub> |  | 2.64% |
| 1783.888 | 0.006 | (Hex)<sub>3</sub> + (Man)<sub>3</sub>(GlcNAc)<sub>2</sub> |  | 4.60% |
| 1794.904 | 0.005 | (Hex)<sub>1</sub> (HexNAc)<sub>1</sub> (Deoxyhexose)<sub>1</sub> + (Man)<sub>3</sub>(GlcNAc)<sub>2</sub> |  | 0.44% |
| 1824.917 | 0.007 | (Hex)<sub>2</sub> (HexNAc)<sub>1</sub> + (Man)<sub>3</sub>(GlcNAc)<sub>2</sub> |  | 0.11% |
| 1987.984 | 0.005 | (Hex)<sub>4</sub> + (Man)<sub>3</sub>(GlcNAc)<sub>2</sub> |  | 3.62% |
| 1999.003 | 0.005 | (Hex)<sub>2</sub> (HexNAc)<sub>1</sub> (Deoxyhexose)<sub>1</sub> + (Man)<sub>3</sub>(GlcNAc)<sub>2</sub> |  | 0.51% |
| 2040.03 | 0.004 | (Hex)<sub>1</sub> (HexNAc)<sub>2</sub> (Deoxyhexose)<sub>1</sub> + (Man)<sub>3</sub>(GlcNAc)<sub>2</sub> |  | 0.28% |
| 2067.041 | 0.004 | (HexNAc)<sub>3</sub> (Pent)<sub>1</sub> + (Man)<sub>3</sub>(GlcNAc)<sub>2</sub> |  | 0.28% |
| 2070.043 | 0.007 | (Hex)<sub>2</sub> (HexNAc)<sub>2</sub> + (Man)<sub>3</sub>(GlcNAc)<sub>2</sub> |  | 0.09% |
| 2081.057 | 0.006 | (HexNAc)<sub>3</sub> (Deoxyhexose)<sub>1</sub> + (Man)<sub>3</sub>(GlcNAc)<sub>2</sub> |  | 4.30% |
| 2192.088 | 0.005 | (Hex)<sub>5</sub> + (Man)<sub>3</sub>(GlcNAc)<sub>2</sub> |  | 5.02% |
| 2244.128 | 0.003 | (Hex)<sub>2</sub> (HexNAc)<sub>2</sub> (Deoxyhexose)<sub>1</sub> + (Man)<sub>3</sub>(GlcNAc)<sub>2</sub> |  | 0.41% |
| 2271.141 | 0.006 | (Hex)<sub>1</sub> (HexNAc)<sub>3</sub> (Pent)<sub>1</sub> + (Man)<sub>3</sub>(GlcNAc)<sub>2</sub> |  | 0.56% |
| 2285.157 | 0.005 | (Hex)<sub>1</sub> (HexNAc)<sub>3</sub> (Deoxyhexose)<sub>1</sub> + (Man)<sub>3</sub>(GlcNAc)<sub>2</sub> |  | 5.74% |
| 2396.187 | 0.004 | (Hex)<sub>6</sub> + (Man)<sub>3</sub>(GlcNAc)<sub>2</sub> |  | 4.95% |
| 2489.258 | 0.006 | (Hex)<sub>2</sub> (HexNAc)<sub>3</sub> (Deoxyhexose)<sub>1</sub> + (Man)<sub>3</sub>(GlcNAc)<sub>2</sub> |  | 1.58% |
| 2600.289 | 0.007 | (Hex)<sub>7</sub> + (Man)<sub>3</sub>(GlcNAc)<sub>2</sub> |  | 0.77% |
| 2652.329 | 0.004 | (Hex)<sub>4</sub> (HexNAc)<sub>2</sub> (Deoxyhexose)<sub>1</sub> + (Man)<sub>3</sub>(GlcNAc)<sub>2</sub> |  | 1.23% |
| 2693.355 | 0.003 | (Hex)<sub>3</sub> (HexNAc)<sub>3</sub> (Deoxyhexose)<sub>1</sub> + (Man)<sub>3</sub>(GlcNAc)<sub>2</sub> |  | 0.08% |
| 2734.389 | 0.012 | (Hex)<sub>2</sub> (HexNAc)<sub>4</sub> (Deoxyhexose)<sub>1</sub> + (Man)<sub>3</sub>(GlcNAc)<sub>2</sub> |  | 0.05% |
| 2897.457 | 0.006 | (Hex)<sub>4</sub> (HexNAc)<sub>3</sub> (Deoxyhexose)<sub>1</sub> +<br>(Man)<sub>3</sub>(GlcNAc)<sub>2</sub> | Multiple forms are possible for this structure | 0.46% |

## References

Aalberse, R. C., Koshte, V., & Clemens, J. G. J. (1981). Immunoglobulin E antibodies that crossreact with vegetable foods, pollen, and Hymenoptera venom. The Journal of Allergy and Clinical Immunology, 68(5), 356–364. doi.org/10.1016/0091-6749(81)90133-0

Adegoke, A., Kumar, D., Bobo, C., Rashid, M. I., Durrani, A. Z., Sajid, M. S., & Karim, S. (2020). Tick-borne pathogens shape the native microbiome within tick vectors. Microorganisms, 8(9), 1–16. doi.org/10.3390/microorganisms8091299

Apostolovic, D., Tran, T. A. T., Hamsten, C., Starkhammar, M., Cirkovic Velickovic, T., & van Hage, M. (2014). Immunoproteomics of processed beef proteins reveal novel galactose-α-1,3-galactose-containing allergens. Allergy, 69(10), 1308–1315. doi.org/10.1111/all.12462

Araujo, R. N., Franco, P. F., Rodrigues, H., Santos, L. C. B., McKay, C. S., Sanhueza, C. A., Brito, C. R. N., Azevedo, M. A., Venuto, A. P., Cowan, P. J., Almeida, I. C., Finn, M. G., & Marques, A. F. (2016). Amblyomma sculptum tick saliva: α-Gal identification, antibody response and possible association with red meat allergy in Brazil. International Journal for Parasitology, 46(3), 213–220. doi.org/10.1016/j.ijpara.2015.12.005

Berg, E. A., Platts-Mills, T. A. E., & Commins, S. P. (2014). Drug allergens and food - The cetuximab and galactose-α-1,3-galactose story. Annals of Allergy, Asthma and Immunology, 112(2), 97–101. doi.org/10.1016/j.anai.2013.11.014

Binder, A. M., Commins, S. P., Altrich, M. L., Wachs, T., Biggerstaff, B. J., Beard, C. B., Petersen, L. R., Kersh, G. J., & Armstrong, P. A. (2021). Diagnostic testing for galactose-alpha-1,3-galactose, United States, 2010 to 2018. Annals of allergy, asthma & immunology : official publication of the American College of Allergy, Asthma, & Immunology, 126(4), 411–416.e1.doi.org/10.1016/j.anai.2020.12.019

Brown, D. B., Muszynski, A., & Carlson, R. W. (2013). Elucidation of a novel lipid A α (1,1)-GalA transferase gene (rgtF) from Mesorhizobium loti: Heterologous expression of rgtF causes Rhizobium etli to synthesize lipid A with α Glycobiology, 23(5), 546–558. doi.org/10.1093/glycob/cws223

Budachetri, K., & Karim, S. (2015). An insight into the functional role of thioredoxin reductase, a selenoprotein, in maintaining normal native microbiota in the Gulf Coast tick (*Amblyomma maculatum*). Insect molecular biology, 24(5), 570–581. https://doi.org/10.1111/imb.12184

Bullard, R. L., Williams, J., & Karim, S. (2016). Temporal gene expression analysis and RNA silencing of single and multiple members of gene family in the lone star tick Amblyomma americanum. PLoS ONE, 11(2), 827–834. doi.org/10.1371/journal.pone.0147966

Bullard, R., Sharma, S. R., Das, P. K., Morgan, S. E., & Karim, S. (2019). Repurposing of Glycine-Rich Proteins in Abiotic and Biotic Stresses in the Lone-Star Tick (Amblyomma americanum). Frontiers in Physiology, 10, 744. doi.org/10.3389/fphys.2019.00744

Cabezas-Cruz, A., Espinosa, P. J., Alberdi, P., Šimo, L., Valdés, J. J., Mateos-Hernández, L., Contreras, M., Rayo, M. V., & de la Fuente, J. (2018). Tick galactosyltransferases are involved in α Anaplasma phagocytophilum infection and Ixodes scapularis tick vector development. Scientific Reports, 8(1), 14224.doi.org/10.1038/s41598-018-32664-z

Cabezas-Cruz, A., Hodžić, A., Román-Carrasco, P., Mateos-Hernández, L., Duscher, G. G., Sinha, D. K., Hemmer, W., Swoboda, I., Estrada-Peña, A., & De La Fuente, J. (2019). Environmental and molecular drivers of the α-Gal syndrome. In Frontiers in Immunology (Vol. 10, Issue MAY, p. 1210). Frontiers Media S.A. doi.org/10.3389/fimmu.2019.01210

Calhoun, D. H., Bishop, D. F., Bernstein, H. S., Quinn, M., Hantzopoulos, P., & Desnick, R. J. (1985). Fabry disease: Isolation of a cDNA clone encoding human *α*-galactosidase A. Proceedings of the National Academy of Sciences of the United States of America, 82(21), 7364–7368. doi.org/10.1073/pnas.82.21.7364

Chai, Y., Beauregard, P. B., Vlamakis, H., Losick, R., & Kolter, R. (2012). Galactose metabolism plays a crucial role in biofilm formation by Bacillus subtilis. MBio, 3(4). doi.org/10.1128/mBio.00184-12

Choudhary, S. K., Karim, S., Iweala, O. I., Choudhary, S., Crispell, G., Sharma, S. R., Addison, C. T., Kulis, M., Herrin, B. H., Little, S. E., & Commins, S. P. (2021). Tick salivary gland extract induces alpha-gal syndrome in alpha-gal deficient mice. Immunity, inflammation and disease, 9(3), 984–990. doi.org/10.1002/iid3.457

Cherepanova, N. A., & Gilmore, R. (2016). Mammalian cells lacking either the cotranslational or posttranslocational oligosaccharyltransferase complex display substrate-dependent defects in asparagine linked glycosylation. Scientific Reports, 6(1), 1–12. doi.org/10.1038/srep20946

Childs, J. E., & Paddock, C. D. (2003). The Ascendancy of Amblyomma americanum as a Vector of Pathogens Affecting Humans in The United States. In Annual Review of Entomology (Vol. 48, pp. 307–337). doi.org/10.1146/annurev.ento.48.091801.112728

Chinuki, Y., Ishiwata, K., Yamaji, K., Takahashi, H., Morita, E. (2016). Haemaphysalis Longicornis Tick Bites Are a Possible Cause of Red Meat Allergy in Japan. Allergy, 71 (3), 421–425. doi: 10.1111/all.12804

Commins, S. P. (2020). Diagnosis & management of alpha-gal syndrome: lessons from 2,500 patients. In Expert Review of Clinical Immunology (pp. 1–11). doi.org/10.1080/1744666X.2020.1782745

Commins, S. P., James, H. R., Kelly, L. A., Pochan, S. L., Workman, L. J., Perzanowski, M. S., Kocan, K. M., Fahy, J. V., Nganga, L. W., Ronmark, E., Cooper, P. J., & Platts-Mills, T. A. E. (2011). The relevance of tick bites to the production of IgE antibodies to the mammalian oligosaccharide galactose-α-1,3-galactose. Journal of Allergy and Clinical Immunology, 127(5), 1286–1293.e6. doi.org/10.1016/j.jaci.2011.02.019

Commins, S. P., & Platts-Mills, T. A. E. (2013). Delayed anaphylaxis to red meat in patients with ige specific for galactose alpha-1,3-galactose (alpha-gal). Current Allergy and Asthma Reports, 13(1), 72–77. doi.org/10.1007/s11882-012-0315-y

Crispell, G., Commins, S. P., Archer-Hartman, S. A., Choudhary, S., Dharmarajan, G., Azadi, P., & Karim, S. (2019). Discovery of alpha-gal-containing antigens in North American tick species believed to induce red meat allergy. Frontiers in Immunology, 10(MAY). doi.org/10.3389/fimmu.2019.01056

Díaz-Alvarez, L., & Ortega, E. (2017). The Many Roles of Galectin-3, a Multifaceted Molecule, in Innate Immune Responses against Pathogens. In Mediators of Inflammation (Vol. 2017).. doi.org/10.1155/2017/9247574

Freeze HH, Hart GW, Schnaar RL. Glycosylation Precursors. 2017. In: Varki A, Cummings RD, Esko JD, et al., editors. Essentials of Glycobiology. 3rd edition. Cold Spring Harbor (NY): Cold Spring Harbor Laboratory Press; 2015-2017. Chapter 5. Available from: https://www.ncbi.nlm.nih.gov/books/NBK453043/ doi: 10.1101/glycobiology.3e.005

Galili, U. (2005). The α-gal epitope and the anti-Gal antibody in xenotransplantation and in cancer immunotherapy. In Immunology and Cell Biology (Vol. 83, Issue 6, pp. 674–686). doi.org/10.1111/j.1440-1711.2005.01366.x

Galili, U. (2015). Significance of the Evolutionary α1,3-Galactosyltransferase (GGTA1) Gene Inactivation in Preventing Extinction of Apes and Old World Monkeys. Journal of Molecular Evolution, 80(1), 1–9. doi.org/10.1007/s00239-014-9652-x

Galili, U., & Avila, J. L. (Eds.). (1999). α-Gal and Anti-Gal, α1,3-Galactosyltransferase, α-Gal Epitopes, and the Natural Anti-Gal Antibody Subcellular Biochemistry Vol. 32 (Boston, MA: Springer US). doi: 10.1007/978-1-4615-4771-6

Goddard, J., & Varela-Stokes, A. S. (2009). Role of the lone star tick, Amblyomma americanum (L.), in human and animal diseases. In Veterinary Parasitology (Vol. 160, Issues 1–2, pp. 1–12). doi.org/10.1016/j.vetpar.2008.10.089

Del Moral, M. G., & Martínez-Naves, E. (2017). The Role of Lipids in Development of Allergic Responses. Immune network, 17(3), 133–143. doi.org/10.4110/in.2017.17.3.133

Hamadeh, R. M., Galili, U., Zhou, P., & Griffiss, J. M. (1995). Anti-α-galactosyl immunoglobulin A (IgA), IgG, and IgM in human secretions. Clinical and Diagnostic Laboratory Immunology, 2(2), 125–131. doi.org/10.1128/cdli.2.2.125-131.1995

Hamsten, C., Tran, T. A. T., Starkhammar, M., Brauner, A., Commins, S. P., Platts-Mills, T. A. E., & van Hage, M. (2013a). Red meat allergy in Sweden: Association with tick sensitization and B-negative blood groups. Journal of Allergy and Clinical Immunology, 132(6), 1431–1434.e6. doi.org/10.1016/j.jaci.2013.07.050

Hamsten, C., Starkhammar, M., Tran, T. A. T., Johansson, M., Bengtsson, U., Ahlén, G., et al. (2013). Identification of Galactose-α-1,3-Galactose in the Gastrointestinal Tract of the Tick Ixodes Ricinus; Possible Relationship With Red Meat Allergy. Allergy, 68 (4), 549–552. doi: 10.1111/all.12128

Hennet, T. (2002). The galactosyltransferase family. In Cellular and Molecular Life Sciences (Vol. 59, Issue 7, pp. 1081–1095). Springer. doi.org/10.1007/s00018-002-8489-4

Hilger, C., Fischer, J., Swiontek, K., Hentges, F., Lehners, C., Eberlein, B., Morisset, M., Biedermann, T., & Ollert, M. (2016). Two galactose-α-1,3-galactose carrying peptidases from pork kidney mediate anaphylactogenic responses in delayed meat allergy. Allergy: European Journal of Allergy and Clinical Immunology, 71(5), 711– 719. doi.org/10.1111/all.12835

Hoxmeier, J. C., Fleshman, A. C., Broeckling, C. D., Prenni, J. E., Dolan, M. C., Gage, K. L., & Eisen, L. (2017). Metabolomics of the tick-Borrelia interaction during the nymphal tick blood meal. Scientific Reports, 7. doi.org/10.1038/srep44394

Huang, X., Tsuji, N., Miyoshi, T., Nakamura-Tsuruta, S., Hirabayashi, J., & Fujisaki, K. (2007). Molecular characterization and oligosaccharide-binding properties of a galectin from the argasid tick Ornithodoros moubata. Glycobiology, 17(3), 313–323. doi.org/10.1093/glycob/cwl070

Iweala, O. I., Choudhary, S. K., Addison, C. T., Batty, C. J., Kapita, C. M., Amelio, C., Schuyler, A. J., Deng, S., Bachelder, E. M., Ainslie, K. M., Savage, P. B., Brennan, P. J., & Commins, S. P. (2020). Glycolipid-mediated basophil activation in alpha-gal allergy. Journal of Allergy and Clinical Immunology, 146(2), 450–452. doi.org/10.1016/j.jaci.2020.02.006

Karim, S., & Ribeiro, J. M. (2015). An Insight into the Sialome of the Lone Star Tick, Amblyomma americanum, with a Glimpse on Its Time Dependent Gene Expression. PloS one, 10(7), e0131292. doi.org/10.1371/journal.pone.0131292

McAuley, M., Kristiansson, H., Huang, M., Pey, A. L., & Timson, D. J. (2016). Galactokinase promiscuity: A question of flexibility? Biochemical Society Transactions, 44(1), 116–122. doi.org/10.1042/BST20150188

Montassier, E., Al-Ghalith, G. A., Mathé, C., Le Bastard, Q., Douillard, V., Garnier, A., Guimon, R., Raimondeau, B., Touchefeu, Y., Duchalais, E., Vince, N., Limou, S., Gourraud, P.-A., Laplaud, D. A., Nicot, A. B., Soulillou, J.-P., & Berthelot, L. (2019). 1,3-Galactosyltransferase Genes in the Human Gut Microbiome. Frontiers in Immunology, 10, 3000. doi.org/10.3389/fimmu.2019.03000

Monzón, J. D., Atkinson, E. G., Henn, B. M., & Benach, J. L. (2016). Population and evolutionary genomics of *Amblyomma americanum*, an expanding arthropod disease vector. Genome Biology and Evolution, 8(5), 1351–1360. doi.org/10.1093/gbe/evw080

Narasimhan, S., Rajeevan, N., Liu, L., Zhao, Y. O., Heisig, J., Pan, J., Eppler-Epstein, R., Deponte, K., Fish, D., & Fikrig, E. (2014). Gut microbiota of the tick vector Ixodes scapularis modulate colonization of the Lyme disease spirochete. Cell Host and Microbe, 15(1), 58–71. doi.org/10.1016/j.chom.2013.12.001

Patrick, C. D., & Hair, J. A. (1975). Laboratory rearing procedures and equipment for multi-host ticks (Acarina: Ixodidae). Journal of Medical Entomology, 12(3), 389– 390.

Raghavan, R. K., Townsend Peterson, A., Cobos, M. E., Ganta, R., & Foley, D. (2019). Current and Future Distribution of the Lone Star Tick, Amblyomma americanum (L.) (Acari: Ixodidae) in North America. PLoS ONE, 14(1). doi.org/10.1371/journal.pone.0209082

Raven, P. H., & Johnson, G. B. (1995). Understanding biology3rd ed. WCB/McGraw- Hill.

Roseman, S. (2001). Reflections on Glycobiology. In Journal of Biological Chemistry (Vol. 276, Issue 45, pp. 41527–41542).. doi.org/10.1074/jbc.R100053200

Sayler, K. A., Barbet, A. F., Chamberlain, C., Clapp, W. L., Alleman, R., Loeb, J. C., Lednicky, J. A., & Kuhn, J. H. (2014). Isolation of tacaribe virus, a caribbean arenavirus, from host-seeking amblyomma americanum ticks in Florida. PLoS ONE, 9(12). doi.org/10.1371/journal.pone.0115769

Sharma SR, Karim S. Tick Saliva and the Alpha-Gal Syndrome: Finding a Needle in a Haystack. Front. Cell. Infect. Microbiol. 11:680264. doi:10.3389/fcimb.2021.680264.

Sicherer, S. H., & Sampson, H. A. (2006). 9. Food allergy. Journal of Allergy and Clinical Immunology, 117(2 SUPPL. 2). doi.org/10.1016/j.jaci.2005.05.048

Sonenshine DE. (2018). Range expansion of tick disease vectors in North America: Implications for spread of tick-borne disease. International Journal of Environmental Research and Public health, 15: 478. doi:10.3390/ijerph15030478

Springer YP, Jarnevich CS, Barnett DT, Monaghan AJ, Eisen RJ. (2015). Modeling the present and future geographic distribution of the lone-star tick Amblyomma americanum (Ixodidae: Ixodida) in the continental United States. Am. J. Trop. Med. Hyg, 93(4):875–890.

Takahashi, H., Chinuki, Y., Tanaka, A., & Morita, E. (2014). Laminin γ-1 and collagen α-1 (VI) chain are galactose-α-1,3-galactose-bound allergens in beef. Allergy: European Journal of Allergy and Clinical Immunology, 69(2), 199–207. doi.org/10.1111/all.12302

Van Nunen, S. A., O’Connor, K. S., Clarke, L. R., Boyle, R. X., & Fernando, S. L. (2009). An association between tick bite reactions and red meat allergy in humans. Medical Journal of Australia, 190(9), 510–511. doi.org/10.5694/j.1326-5377.2009.tb02533.x

Wong, L. J., & Frey, P. A. (1974). Galactose-1-Phosphate Uridylyltransferase. Rate Studies Confirming a Uridylyl-Enzyme Intermediate on the Catalytic Pathway. Biochemistry, 13(19), 3889–3894. doi.org/10.1021/bi00716a011

Wuerdeman, M. F., & Harrison, J. M. (2014). A case of tick-bite-induced red meat allergy. Military Medicine, 179(4), e473–e475. doi.org/10.7205/MILMED-D-13-00369

